# Stabilization of the trimeric pre-fusion structures of influenza H1 and H9 hemagglutinins by mutations in the stem helices

**DOI:** 10.1101/2025.08.27.672506

**Authors:** Ga Young Lee, Yun Geun Kim, Hee-Su Shin, Chang Jun Lee, Jie-Oh Lee

**Author notes:** These authors contributed equally: Ga Young Lee, Yun Geun Kim.

## Abstract

Stabilizing the pre-fusion structures of antigenic proteins can enhance the effectiveness of antiviral vaccines. The pre-fusion form of hemagglutinin (HA) from the influenza virus typically adopts a stable trimeric structure. However, the recombinant ectodomain of HA from the A/California/04/2009 (H1N1) influenza virus formed a monomer in solution rather than the expected trimer. To promote trimer formation in the pre-fusion conformation, we redesigned five amino acid residues in the stem region of HA that are involved in trimerization. The engineered HA protein formed a stable trimer at both pH 8.0 and pH 5.5. Additionally, the thermal stability of the modified protein improved, as indicated by an approximately ten-degree increase in its denaturation temperature. Cryo-EM analysis at 2.2 Å resolution confirmed that the mutant HA protein adopted the pre-fusion structure. Furthermore, the stabilized mutant exhibited enhanced immunogenicity in mice. We applied the same optimization strategy to the HA proteins from A/Malaysia/1706215/2007 (H1N1) and A/swine/Hong Kong/2106/98 (H9N2). These engineered proteins demonstrated increased thermal stability and retained a trimeric pre-fusion structure, as confirmed by cryo-EM analysis. Extending this optimization strategy to the equivalent five residues in hemagglutinins from six additional group 1 influenza viruses successfully stabilized their trimeric structures.

## Introduction

Influenza remains a persistent threat to global public health, with periodic epidemics and pandemics causing significant morbidity and mortality. Central to the influenza virus’s ability to infect host cells is the hemagglutinin (HA) protein, which is crucial for viral entry and is the primary target for neutralizing antibodies ^1^. HA facilitates virus binding to sialic acid receptors on the host cell surface, promoting fusion of the viral membrane with the host cell membrane, thus enabling viral entry. This fusion process depends on a conformational change in HA from a pre-fusion to a post-fusion state ^2–5^. Consequently, stabilizing HA in its pre-fusion form is critical for vaccine development, as this form is the most effective target for eliciting protective immune responses. Stabilizing the pre-fusion structure has proven effective in the development of vaccines against RSV and COVID-19 ^6–9^.

The HA protein is synthesized as a precursor polypeptide, HA0, which is cleaved into two subunits, HA1 and HA2, linked by disulfide bonds ^10^. HA1 contains the receptor-binding domain, while HA2 facilitates membrane fusion. Structurally, HA is expressed on the viral surface as a trimer, with each monomer composed of one HA1 and one HA2 subunit. This trimeric arrangement is essential for HA function, enabling efficient binding to sialic acid receptors on host cells, leading to viral entry through endocytosis. Acidification within the endosome triggers a conformational change in HA, exposing the fusion peptide located in the HA2 subunit, thereby facilitating membrane fusion ^11–14^. Stabilizing the pre-fusion structure is beneficial for vaccine development, as the pre-fusion form is critical for recognition by most neutralizing antibodies ^10^.

Despite the critical role of the pre-fusion form of HA in vaccine development, stabilizing it poses significant challenges. To overcome these challenges, various strategies have been employed. One common approach involves incorporating foreign trimerization domains, such as the T4 Foldon or ferritin cages, to promote trimer formation. For example, Weldon *et al.* demonstrated that incorporating a trimerization sequence tag into the H3N2 HA protein allowed the engineered HA to form native-like trimers, resulting in enhanced vaccine activity in mice ^15^. In the case of the 24-subunit ferritin cage, monomeric ferritin subunits assemble into trimers at the vertices. When HA monomers were attached to the ferritin cage, a stable trimeric arrangement formed, providing potent and broad protection against diverse influenza strains ^16^. Ferritin-based HA vaccines have shown promising results in preclinical and early clinical trials, exhibiting strong immunogenicity and protective efficacy ^17^. Additionally, an isolated HA stem protein, stabilized by N- and C-terminal trimerization domains and engineered disulfide bridges demonstrated broad protection against various influenza viruses ^18^.

Here, we investigated an efficient strategy to stabilize the pre-fusion state of HA. Our research demonstrates that optimizing the amino acid sequence at a limited number of pre-selected positions in HA, using a rapid computational method, effectively stabilizes the protein in its trimeric pre-fusion form. This strategy not only enhances thermal stability but also preserves the pre-fusion structure of HA, as confirmed by cryo-EM analysis. Identifying multiple sets of stabilizing amino acid substitutions and validating them experimentally in advance will improve our preparedness for future influenza pandemics. This is important because relying on only one or two stabilization strategies may be insufficient against emerging influenza strains with unexpected sequence variations.

## Results

### Wild-type A/California/04/2009 (H1N1) HA ectodomain does not form the trimeric pre-fusion structure

To determine the structure of HA, we produced the wild type ectodomain of H1 HA from the influenza strain A/California/04/2009 (H1N1), H1/Cal09, with Genbank id ACP41105 (Fig. 1a). Unexpectedly, the purified protein eluted as a monomer during gel permeation chromatography, indicating that the stable trimeric pre-fusion structure may not form (Fig. 1b). 2D classification analysis using cryo-EM confirmed that the ectodomain of H1/Cal09 HA predominantly exists as a monomer (Fig. 1c). To enhance the formation of the trimeric pre-fusion structure, we attached the Foldon trimerization sequence to the C-terminus of the HA ectodomain (Fig. 2a). It is a short sequence composed of 12 amino acids derived from the trimerization domain of T4 fibritin ^19^. The Foldon can induce trimerization of the host protein due to its unusually stable trimeric structure and fast folding kinetics. This modification to the HA ectodomain shifted the gel permeation chromatographic elution profile of the protein from a monomer to a trimer as anticipated (Fig. 2b). However, upon collecting the trimeric fractions and inspecting their structure using cryo-EM, we found that the protein is indeed a trimer but does not have a stable structure, and high-resolution 3D reconstruction of the structure was impossible (Fig. 2c). This analysis demonstrates that trimerization induced by the Foldon sequence tag does not ensure the formation of the pre-fusion structure of the H1/Cal09 HA protein.

**Fig. 1.**
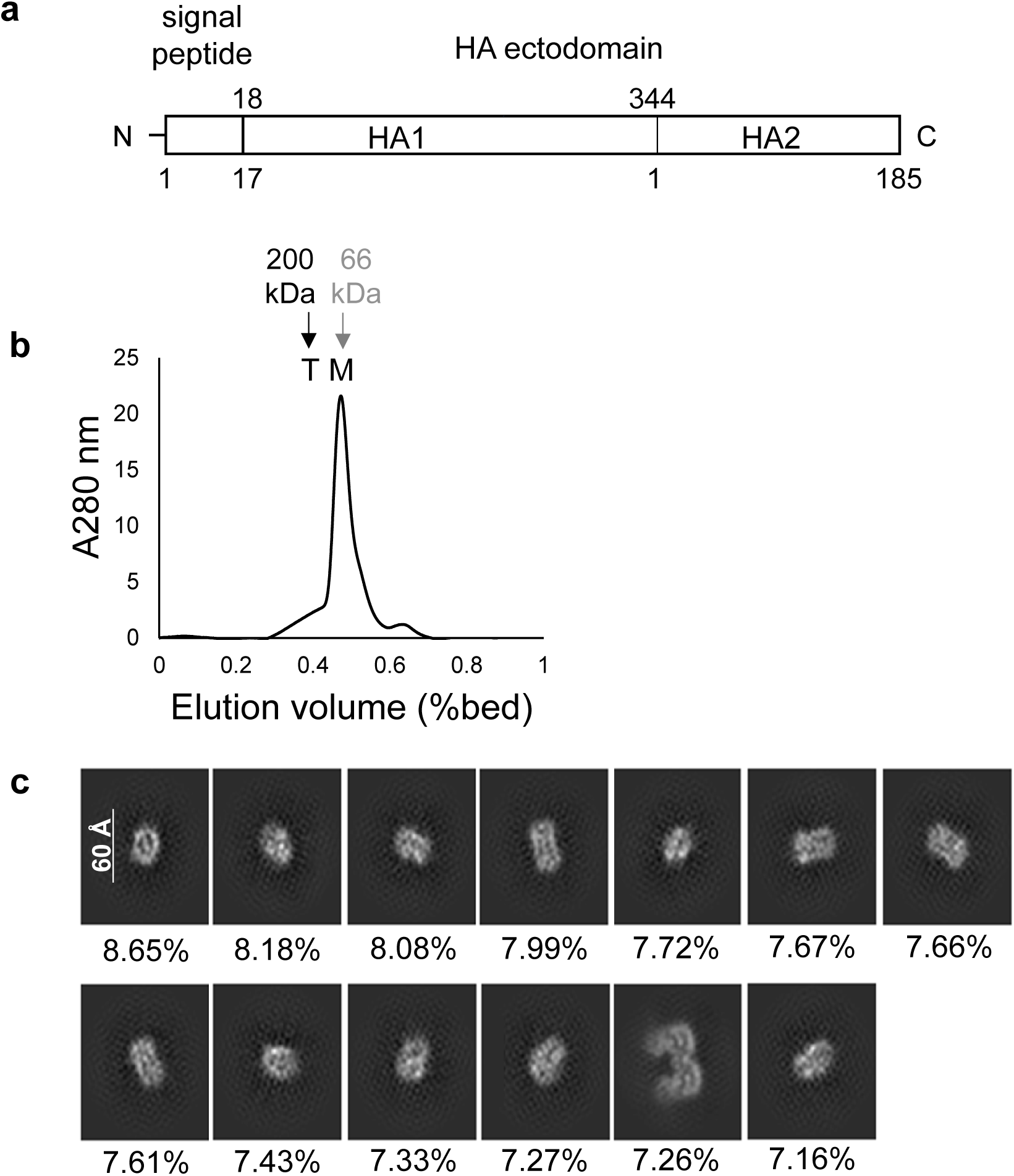
Trimerization state of the ectodomain of HA from the A/California/04/2009 (H1N1) virus. **a** Domain architecture of the HA ectodomain. **b** Chromatographic elution profile of the A/California/04/2009 (H1N1) (H1/Cal09) HA ectodomain. The purified protein was analyzed using Superdex 200 gel permeation chromatography. The expected elution positions of the peaks for the trimeric and monomeric proteins are indicated by arrows and labeled T and M, respectively. Elution volumes of β-amylase (200 kDa) and BSA (66 kDa) are indicated by black and gray arrows, respectively. **c** 2D class averages from cryo-EM images of the H1/Cal09 HA ectodomain. The relative abundance of each structural class is indicated below each class. The majority of particles correspond to the monomeric form, while a single class, comprising 7.26% of the particles, appears to represent a dimeric form.

**Fig. 2.**
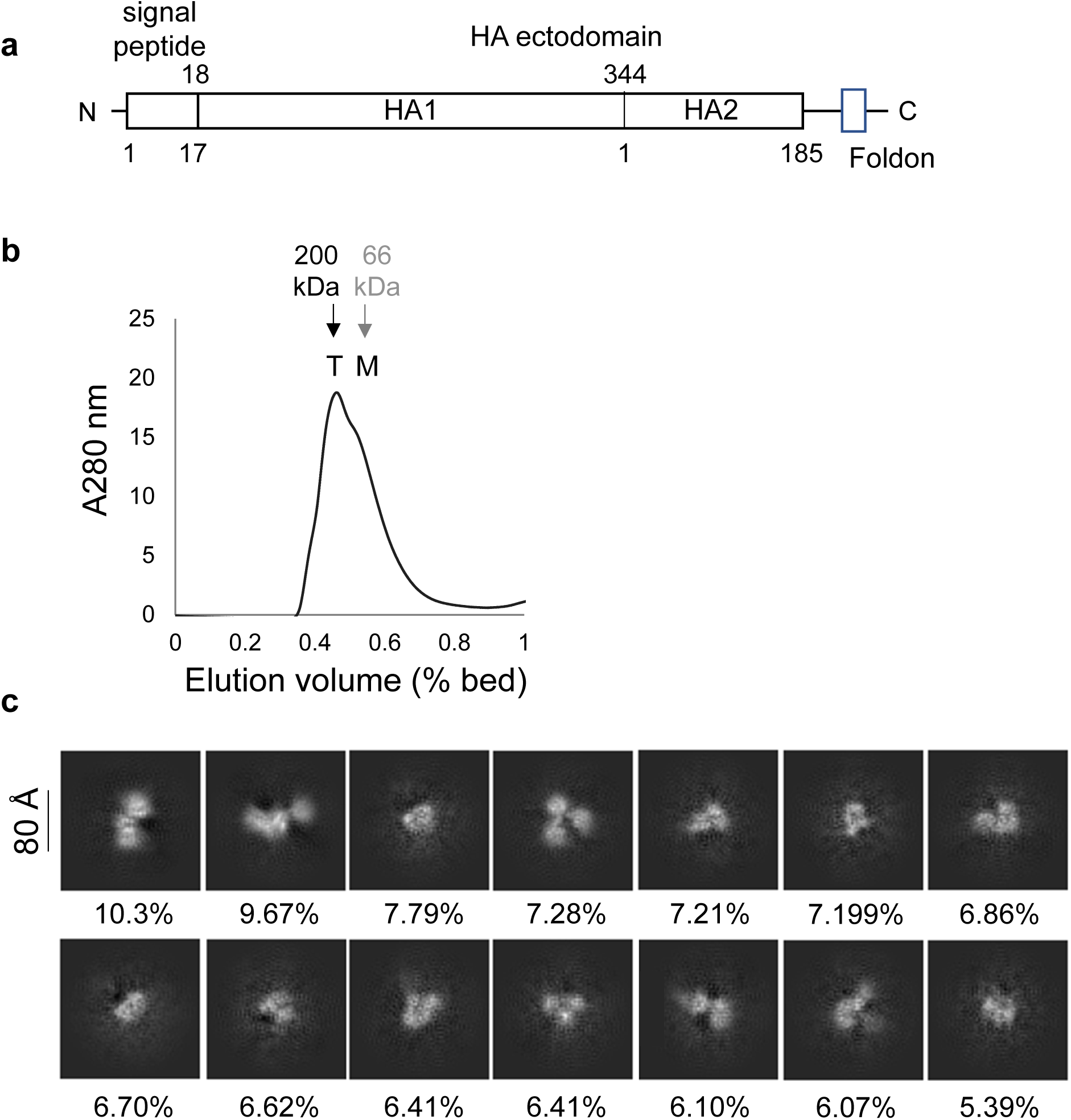
Trimerization state of the Foldon-tagged ectodomain of the HA protein from the H1/Cal09 virus. **a** Domain architecture of the HA ectodomain. The Foldon trimerization domain from T4 fibritin is fused to the C-terminus of the HA ectodomain. **b** Chromatographic elution profile of the Foldon-tagged H1/Cal09 HA ectodomain. The purified protein was analyzed by Superdex 200 gel filtration chromatography. The expected elution positions of the trimeric and monomeric forms are indicated by arrows and labeled T and M, respectively. Elution volumes of β-amylase (200 kDa) and BSA (66 kDa) are indicated by black and gray arrows, respectively. **c** 2D class averages from cryo-EM images of the Foldon-tagged H1/Cal09 HA ectodomain. The relative abundance of each 2D class average is shown below each class. Most particle images correspond to the trimeric form; however, high-resolution 3D structure reconstruction was not possible, presumably due to flexibility in the relative orientations of the monomeric subunits within the trimer.

### Selection of mutational sites and sequence optimization through computational design

To stabilize the trimeric pre-fusion structure of the H1/Cal09 HA with a minimal number of mutations, we examined the trimerization interface formed by the central stem alpha helices in the previously reported structure with PDB id 7MEM (Supplementary Fig. 1). We identified the following five amino acid residues that did not appear optimal for stable trimerization of the HA monomers (Supplementary Fig. 2). The negatively charged E47 of the HA2 chain is positioned near the hydrophobic L30 of a neighboring subunit. Hydrophilic N95 is located at the center of the hydrophobic core of the trimerization interface, surrounded by I91, W92, Y94, L98, and L99. Additionally, hydrophobic L102 is flanked by charged residues E103 and R106, while charged E103 is in contact with the hydrophobic L102. Finally, D109 is situated at the periphery of the trimerization interface. It appears that longer amino acids at this position may enhance interaction with the other two subunits in the trimerization interface. Among these, N95, L102, E103, and D109 residues are located in the central stem alpha helix and are directly involved in the trimerization. E47 is not in the trimerization interface but mediates interaction with neighboring HA monomers and, therefore, is part of the dimerization interface between the two interacting HA monomers.

These five amino acids underwent computational optimization using the protein design program protein MPNN ^20^. The previously reported structure of H1/Cal09 HA was used as the structural template for the calculation ^21^. This deep learning-based program calculates the amino acid sequences that fit best to the backbone structure provided as a template. Following a quick optimization run, E47 was replaced with Gly, N95 with Leu, L102 with Phe, E103 with Leu, and D109 with Glu, resulting in a substantial decrease in the computational score value. In the context of protein MPNN, a lower score indicates a more stable mutation associated with lower free energy.

### Stabilization of pre-fusion structure of the H1/Cal09 HA protein

To test the stabilization effect of the computational sequence optimization, we produced various proteins: the wild-type ectodomain (H1/Cal09-WT), the wild-type with the Foldon trimerization tag (H1/Cal09-WT(tri)), and a mutant with five mutations (H1/Cal09-mut). These proteins were analyzed using gel permeation chromatography to assess their trimerization state at three different pH levels (Fig. 3a). At both pH 8.0 and 5.5, the wild-type HA eluted as a monomer, while the H1/Cal09-WT(tri), and the five-mutant variant, H1/Cal09-mut, eluted as trimers. At pH 4.7, all three proteins appeared unstable, with less than half of the protein injected into the column eluted, suggesting that a substantial portion aggregated in the column and did not elute.

**Fig. 3.**
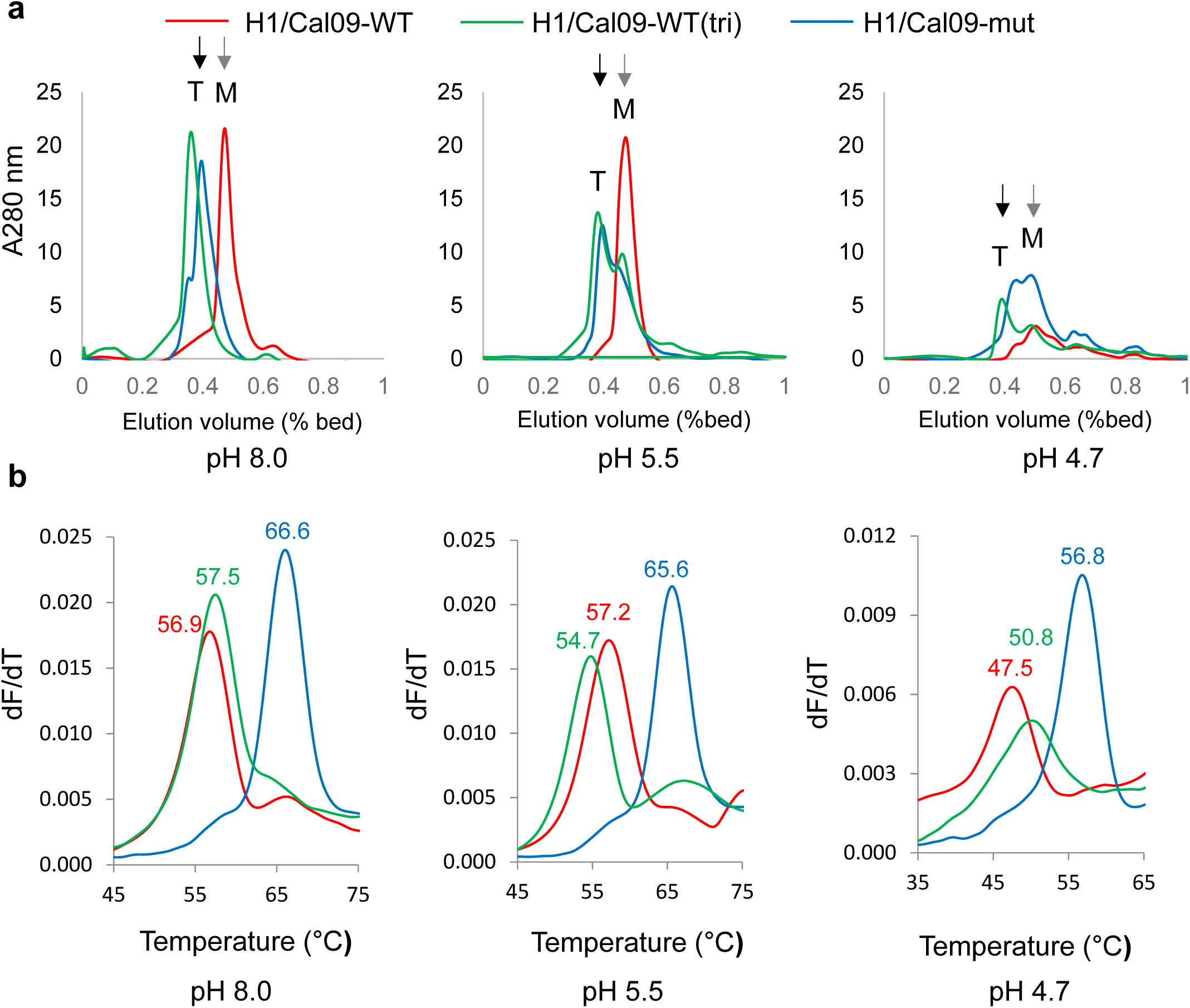
Trimerization state and thermal stability of H1/Cal09 HA ectodomain wild-type and mutant proteins. **a** Gel permeation chromatographic elution profiles of the wild-type, wild-type with Foldon tag, mutant, and mutant with Foldon tag proteins, analyzed at three different pH values. Elution volumes of β-amylase (200 kDa) and BSA (66 kDa) are indicated by black and gray arrows, respectively. **b** Differential scanning fluorimetry (DSF) profiles of the wild-type and mutant HA ectodomain proteins.

To assess the thermal stability of the wild-type and mutant proteins, we measured their unfolding temperature, Tm, using differential scanning fluorometry (DSF) (Fig. 3b). This method monitors protein unfolding through intrinsic tryptophan fluorescence. At all three pH levels, the Foldon sequence tag stabilized only 1∼3 degrees on the thermal stability. However, our five-site mutation enhanced the stability of the HA proteins by ∼10 degrees. These data demonstrate that mutations at the trimerization interface stabilize the protein structure and promote trimerization. The structure of the five-mutation HA protein was analyzed using high-resolution cryo-EM and compared to the pre-fusion structure of H1/Cal09 previously determined with the PDB id, 7MEM. In the 7MEM structure, bound antibodies appear to stabilize the pre-fusion conformation of the HA protein. Our cryo-EM structure of the stabilized mutant, resolved at 2.2 Å, closely resembles the pre-fusion structure of the wild-type HA even in the absence of bound antibodies (Fig. 4a and b). Close-up views of the mutated sites demonstrate that the mutations did not disturb the overall structure of the protein; the side chains of the mutated residues fit seamlessly into the structure, requiring only minor adjustments (Fig. 4c, d and e). In conclusion, our five-site mutation strategy effectively stabilizes the pre-fusion structure of the HA protein of H1/Cal09.

**Fig. 4.**
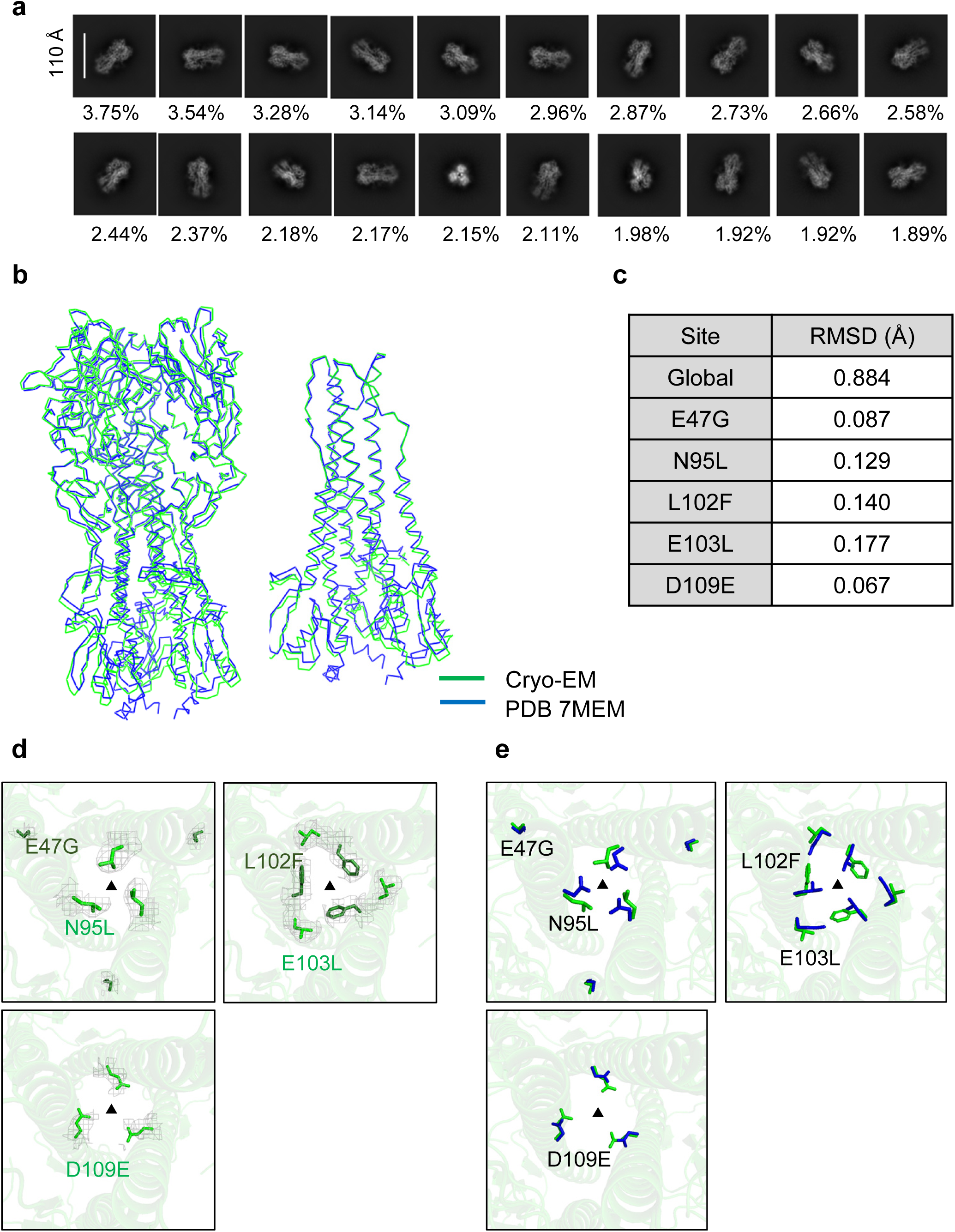
Cryo-EM structure of the mutated H1/Cal09 HA ectodomain. **a** 2D class averages of cryo-EM images of the mutated HA ectodomain. The relative abundance of each structural class is indicated below. **b** Structural comparison of the mutated and wild-type HA. Cα traces of the mutated HA ectodomain (green) and the wild-type HA (PDB ID: 7MEM, blue) are shown. The left panel displays the overall structure of the entire HA ectodomain, while the right panel presents a close-up view of the stem region. **c** RMSD table of the mutated HA ectodomain relative to the wild-type. RMSD values were calculated for five residues neighboring each mutation site to compare the wild-type and mutant HA proteins. **d** Close-up view of the mutated sites. The electron density map around each mutated residue is shown as a gray mesh. The view is from above the stem helices of the trimeric HA, highlighting all three monomeric subunits. The E47G and L102F mutated residues are shown in dark green. All other mutated residues are shown in light green. The electron density map surrounding each mutated residue is displayed as a gray mesh. The threefold symmetry axis is marked by a triangle. **e** Additional close-up view of the mutated sites. Side chains in the wild-type structure (PDB ID: 7MEM) are shown in blue, while those in the cryo-EM structure of the mutant are shown in green.

To test the effect of the stabilizing mutation on the immunogenicity of the protein, we injected wild-type and mutant HA proteins into mice. The proteins were administered intramuscularly, followed by two booster injections to enhance the immune response. Antibody levels in the blood were subsequently measured using the ELISA method (Fig. 5a). As shown in Fig. 5b and c, antibody levels increased approximately 5-fold and 7-fold when wild-type and mutant proteins, respectively, were used as the coating antigens. These results demonstrate that the pre-fusion stabilized H1/Cal09 HA protein elicits a stronger immune response compared to the monomeric wild-type HA protein.

**Fig. 5.**
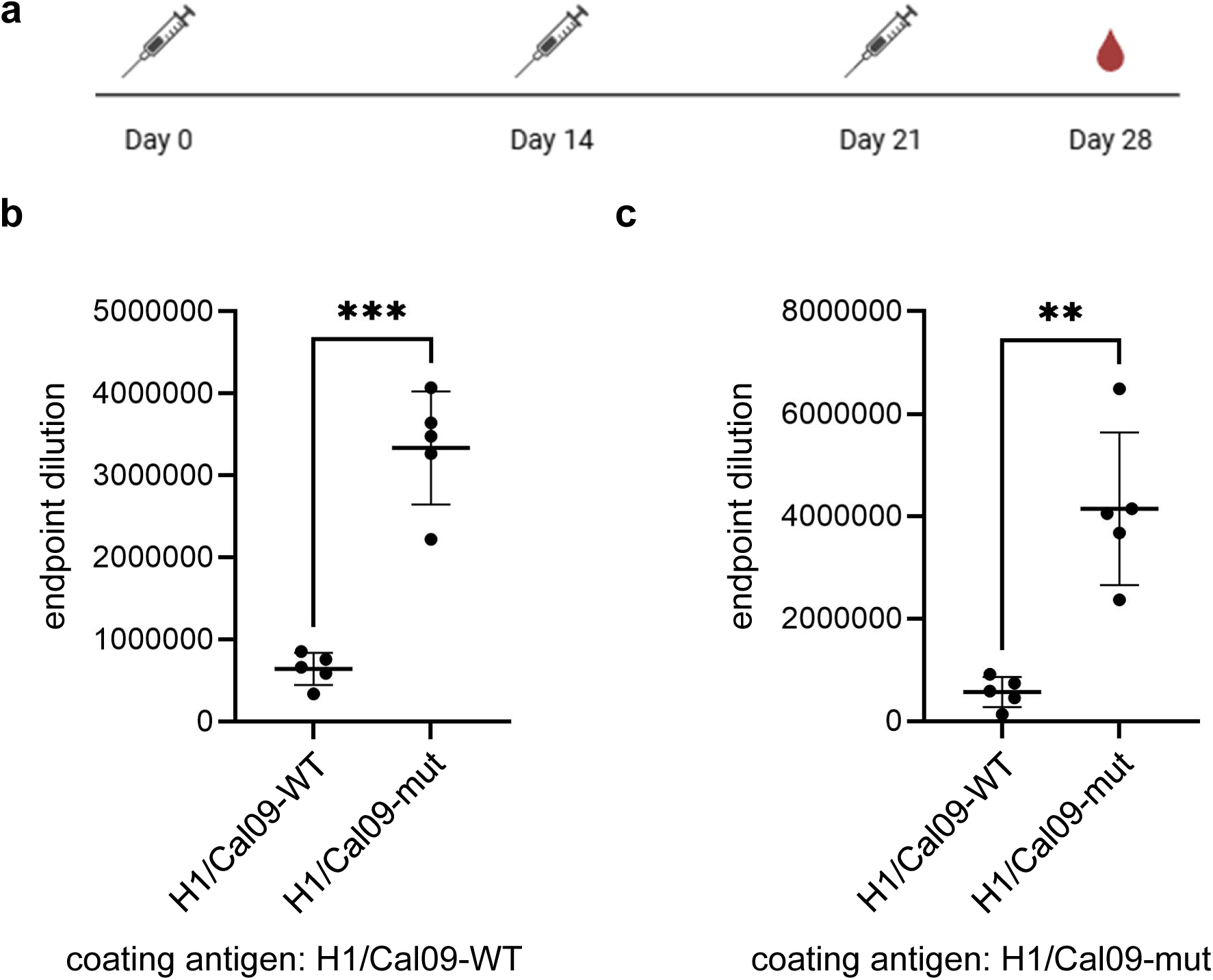
Serum IgG responses induced by wild-type and mutant H1/Cal09 HA ectodomains. **a** Female BALB/c mice (n = 5 per group) were immunized intramuscularly with either wild-type or mutant H1/Cal09 HA protein formulated with Freund’s adjuvant. A second dose was administered two weeks later, followed by a third dose one week after the second. Serum samples were collected one week after the final immunization via retro-orbital bleeding. **b** IgG endpoint titers were measured by ELISA using wild-type H1/Cal09 HA protein as the coating antigen. **c** IgG endpoint titers were measured by ELISA using mutant H1/Cal09 HA protein as the coating antigen. Endpoint titers were defined as the highest serum dilutions yielding OD₄₅₀ values at least twofold greater than those of naïve mice at the lowest dilution (1:100). Each dot represents an individual mouse; bars indicate the mean ± SD. Statistical significance was determined using Welch’s t-test. **P < 0.01; ***P < 0.001.

### Stabilization of the pre-fusion structure of the A/Malaysia/1706215/2007 (H1N1) HA protein

To determine if our five-site mutation strategy is applicable to other H1 HA proteins, we applied it to the A/Malaysia/1706215/2007 (H1N1) (H1/Mal07) HA protein. The HA protein sequences from H1/Cal09 and H1/Mal07 exhibit 79% sequence identity (Supplementary Fig. 3). Sequence variations are primarily clustered in the head region, while the stem regions are highly conserved. Among the five sites we selected for mutation, four, N95, L102, E103, and D109, are conserved in the H1/Mal07 HA protein. Interestingly, the remaining site, E47, is already altered to glycine in the wild type H1/Mal07 HA variant even without the computation optimization. Therefore, only four of the five sites were selected for mutation. We produced the wild-type HA protein (H1/Mal07-WT), the wild-type protein with the Foldon tag (H1/Mal07-WT(tri)), the mutated protein (H1/Mal07-mut), and the mutated protein with the Foldon tag (H1/Mal07-mut(tri)). Their trimerization states were assessed using gel permeation chromatography. As shown in Fig. 6a, the wild-type ectodomain of the H1/Mal07 HA protein eluted as a mixture of trimer and monomer. The addition of the Foldon tag facilitated the trimerization of the wild-type protein. Meanwhile, the mutated protein without the Foldon tag was eluted as a trimer, indicating that the mutations successfully stabilized the trimeric structure even without the Foldon tag. At lower pH levels, the peaks for the wild-type and wild-type Foldon-tagged proteins diminished, presumably due to aggregation, whereas the mutated proteins largely maintained their trimeric state.

**Fig. 6.**
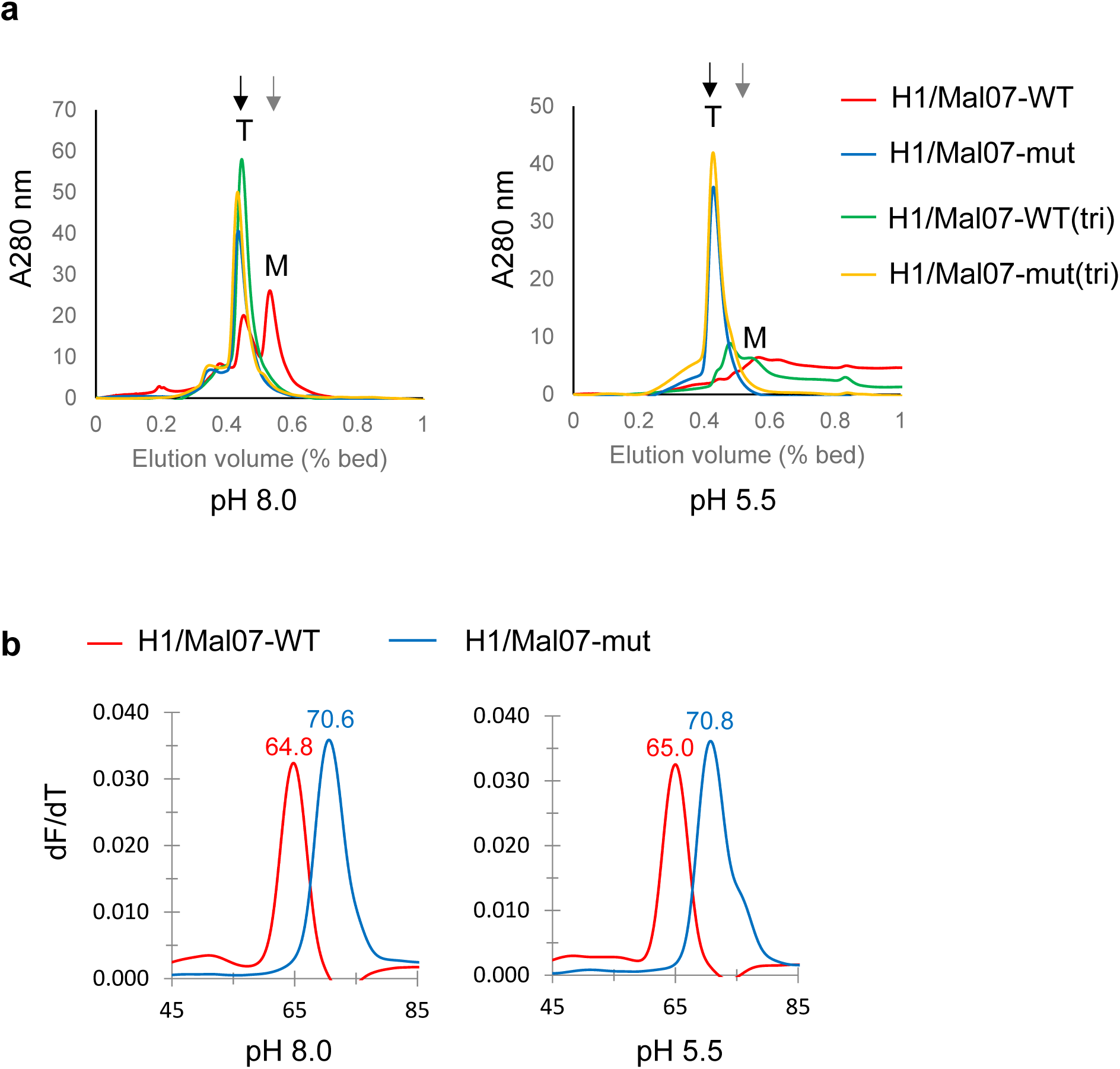
Trimerization state and temperature stability of A/Malaysia/1706215/2007 (H1N1) (H1/Mal07) HA ectodomain wild type and mutants. **a** Gel permeation chromatography elution profiles of the wild type, the wild type with the Foldon tag, the mutant, and the mutant with the Foldon tag proteins at pH 8.0 and 5.5. Elution volumes of β-amylase (200 kDa) and BSA (66 kDa) are indicated by black and gray arrows, respectively. **b** The differential scanning fluorometry (DSF) profiles of these proteins are shown below.

The thermal stability of the mutated protein was assessed using the DSF method (Fig. 6b). At both pH 5.5 and 8.0, the mutated protein exhibited 5.8-degree higher Tm values, demonstrating successful stabilization of the protein structure through the mutation. The structure of the mutated H1/Mal07 HA protein without the Foldon tag was analyzed using cryo-EM at 3.0 Å resolution. As shown in Fig. 7a, the 2D class-averaged maps clearly indicate a trimeric pre-fusion structure. The refined structure was superimposable with that of the reported pre-fusion structure with PDB id of 7UMM (Fig. 7b). The Cα r.m.s.d. value of the two structures was 0.865 Å. Furthermore, a close-up view near the mutated sites shows that the overall structure of the protein was not disturbed by the mutations (Fig. 7c, d and e).

**Fig. 7.**
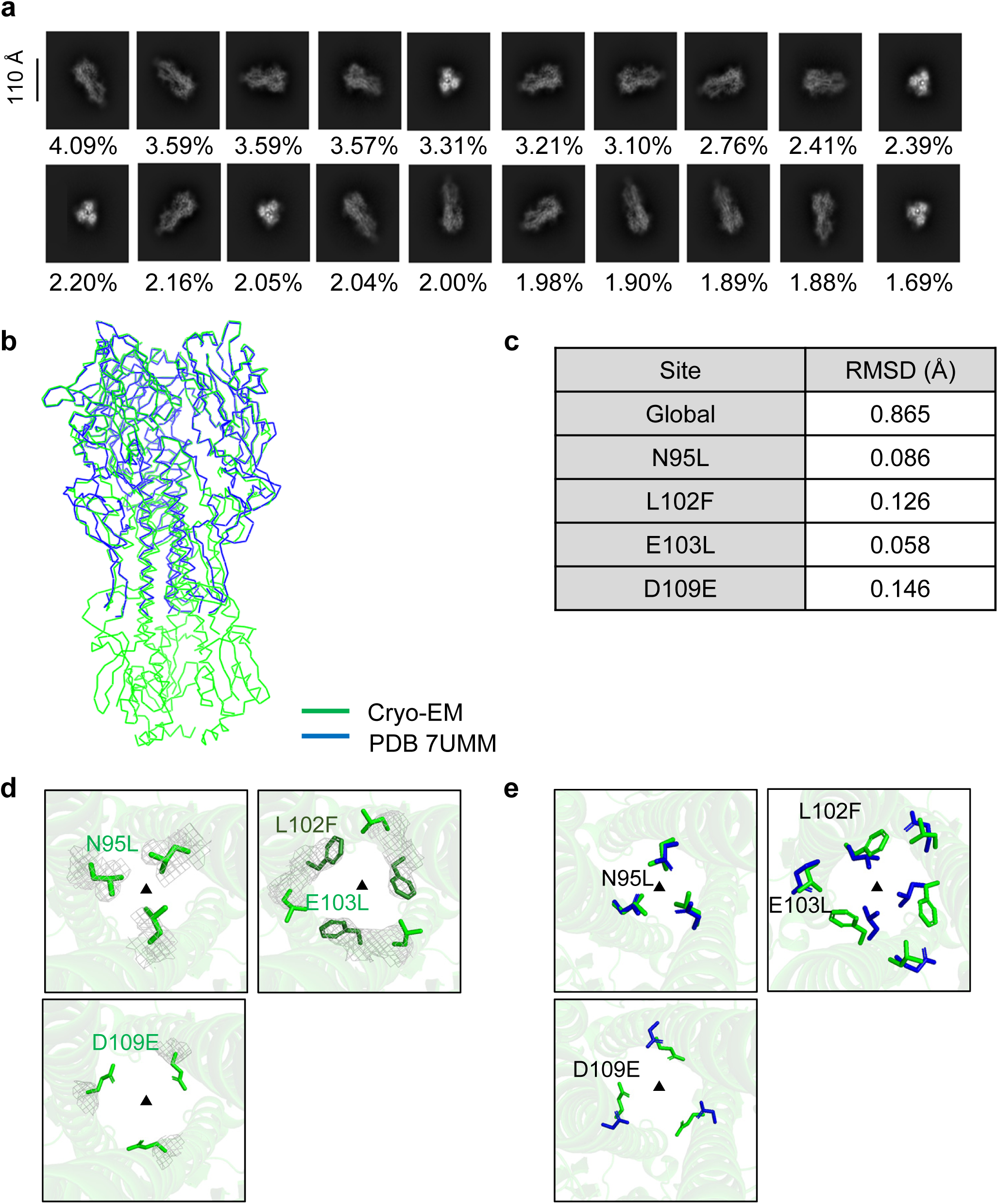
Cryo-EM structure of the mutated H1/Mal07 HA ectodomain. **a** 2D class averages from cryo-EM images of the mutated HA ectodomain. The relative population of each structural class is indicated below. **b** Structural comparison of the mutated and wild-type HA. Cα traces of the mutated (green) and the wild-type HA (PDB ID: 7UMM, blue) are shown. The previously reported crystal structure of the wild-type protein includes only a portion of the stem region. **c** RMSD table of the mutated HA ectodomain relative to the wild-type. RMSD values were calculated for five residues neighboring each mutation site to compare the wild-type and mutant HA proteins. **d** Close-up view of the mutated sites. The electron density map around each mutated residue is shown as a gray mesh. The view is from above the stem helices of the trimeric HA, highlighting all three monomeric subunits. The three L102F mutated residues are shown in dark green. The E103L and all other mutated residues are shown in light green. The electron density map surrounding each mutated residue is displayed as a gray mesh. The threefold symmetry axis is marked by a triangle. **e** Structural comparison of the mutated and wild-type residues. Residues from the wild-type structure (PDB ID: 7UMM) are shown in blue, and those from the mutant structure are shown in green.

To assess the stabilizing role of each mutated amino acid, we individually reverted each residue to its wild-type counterpart and evaluated the oligomerization state and thermal stability of the resulting proteins at both pH 8.0 and pH 5.5 (Fig. 8 and Supplementary Fig. 4). The original mutant protein, H1/Mal07-mut, which contains four mutations, E47G, N95L, L102F, and D109E, was used as the reference. All four reverted mutants formed trimers, as confirmed by gel-permeation chromatography. In the thermal stability assay, the mutant in which leucine at position 95 was reverted to asparagine (designated rev-95L) exhibited a ∼3°C decrease in the denaturation temperature compared to the reference mutant. This indicates that the N95L mutation plays the most important role in stabilizing the prefusion state, contributing approximately 2.8°C of stabilization. The combination of the remaining three mutations, E47G, L102F, and D109E, accounted for the remaining ∼3°C of stabilization. Individual reversion of these three mutations did not result in a noticeable change in denaturation temperature, suggesting that their stabilizing effects are cooperative and that any two of them are sufficient to maintain stability. Similar results were observed at pH 5.5.

**Fig. 8.**
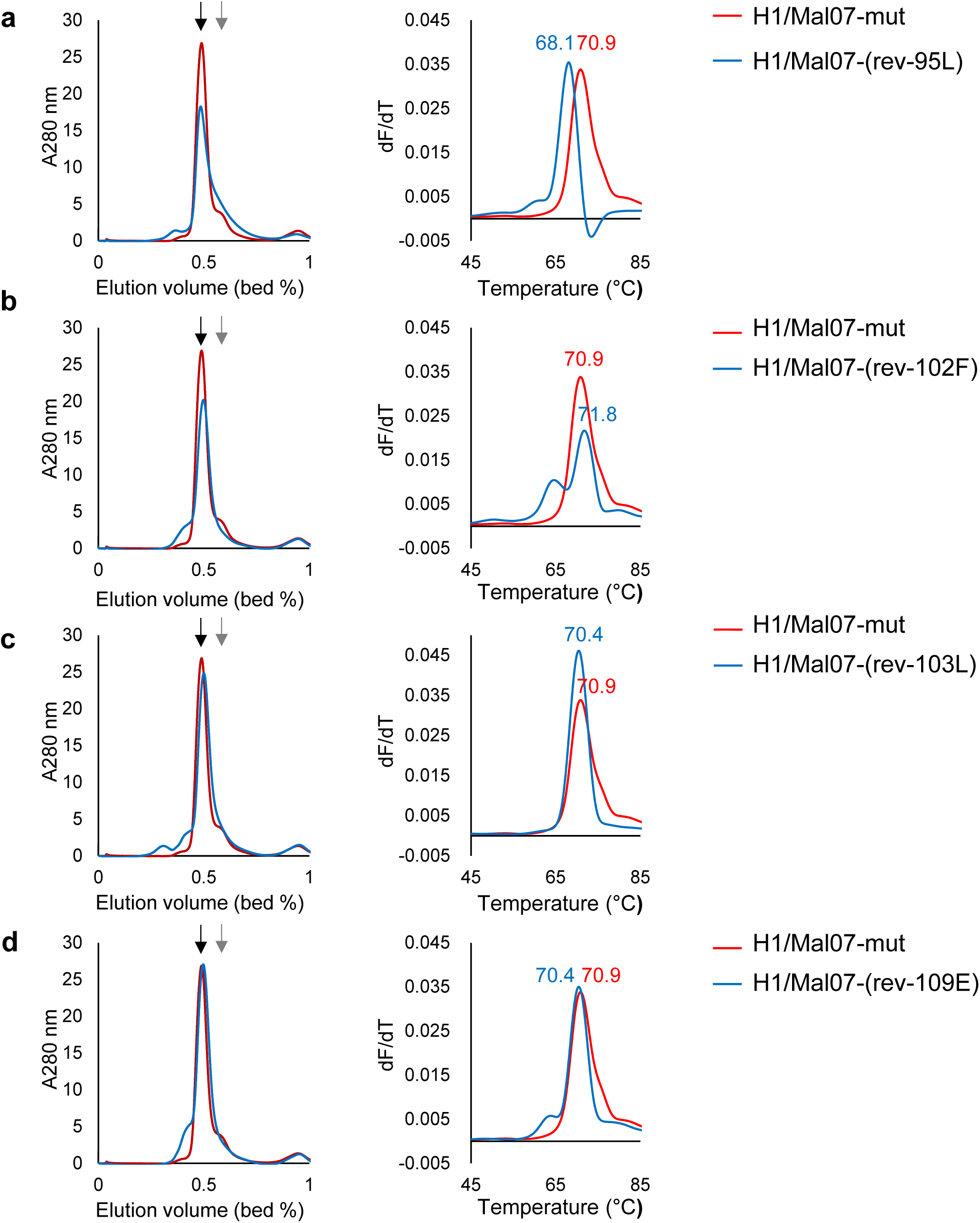
Trimerization state and thermal stability at pH 8.0 of wild-type and mutant H1/Mal07 HA ectodomains. **a-d** Gel permeation chromatography (left) and differential scanning fluorimetry (DSF) profiles (right) of H1/Mal07 HA containing all four mutations (H1/Mal07-mut) or containing only three of the four mutations, with one mutation reverted back to the wild-type sequence. All experiments were conducted at pH 8.0. Elution volumes of β-amylase (200 kDa) and BSA (66 kDa) are indicated by black and gray arrows, respectively.

### Stabilization of pre-fusion structure of the A/swine/Hong Kong/2106/98 (H9N2) HA protein

To assess whether our 5-site optimization strategy is applicable to proteins other than H1 HA proteins, we applied it to the HA protein from an H9 influenza virus, A/swine/Hong Kong/2106/98 (H9/N2), H9/HK98. The sequences of the HA proteins from H1/Cal09 and H9/HK98 were aligned, revealing a 49% sequence identity (Supplementary Fig. 5). Variations in sequences were observed in both the head and stem regions, although the stem area has lower variation. Among the five sites we selected, four, N95, L102, E103, and D109, are conserved in H9/HK98. The remaining E47 residue is altered to lysine in the H9 HA. These five sites were subjected to the computational optimization method using the protein MPNN program. The previously reported structure with PDB id, 1JSD was used as the structural template ^22^.

Unlike H1/Cal09, only two of the five sites were changed by the program (Fig. 9a). We produced the wild type, wild type with the Foldon tag, mutated protein, and mutated protein with the Foldon tag attached. Their trimerization states were analyzed using gel permeation chromatography. As depicted in Fig. 9b, the wild type ectodomain of the H9 HA protein eluted as a mixture of trimer and monomer. The addition of the Foldon tag promoted trimerization in the wild-type protein. The mutated protein without the Foldon tag eluted as a trimer, demonstrating that the mutation successfully stabilized the trimeric structure. At lower pH levels, the peaks for both the wild-type and wild-type Foldon proteins diminished, presumably due to aggregation. In contrast, the mutated proteins largely maintained their trimeric state, suggesting that the mutation enhanced resistance to pH shock.

**Fig. 9.**
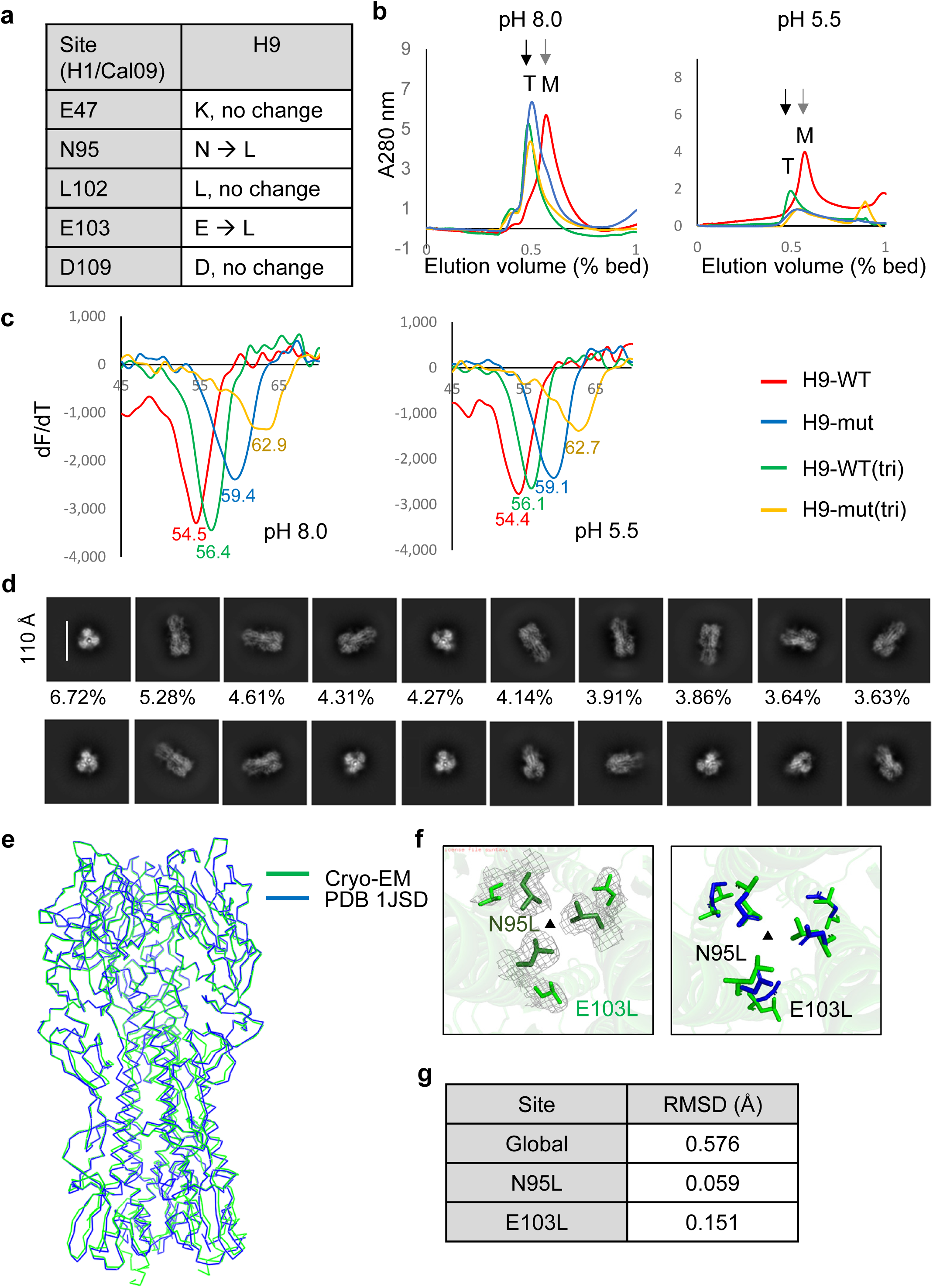
Trimerization state, thermal stability, and cryo-EM structure of the mutated A/swine/Hong Kong/2106/98 (H9N2) (H9/HK98) HA ectodomain. **a** Summary of the stabilizing mutations. **b** Gel permeation chromatographic elution profiles of the wild-type and mutant proteins at three different pH values. Elution volumes of β-amylase (200 kDa) and BSA (66 kDa) are indicated by black and gray arrows, respectively. **c** Differential scanning fluorimetry (DSF) profiles of the wild-type and mutant ectodomains at three different pH values. Denaturation temperatures (Tm) are indicated. **d** 2D class averages from cryo-EM images of the mutated HA ectodomain. **e** Structural comparison of the mutated and wild-type HA. Cα traces of the mutated HA (green) and the wild-type HA (PDB ID: 1JSD, blue) are shown. **f** Close-up view of the mutated sites. (left) Close-up view of the mutated sites. The three N95L mutated residues are shown in dark green, and the E103L mutated residues are shown in light green. The electron density map surrounding each mutated residue is displayed as a gray mesh. (Right) Structural comparison of the mutated and wild-type residues. Residues from the wild-type structure (PDB ID: 1JSD) are shown in blue, and those from the mutant structure are shown in green. The top-down view displays the stem helices of the three HA subunits, with the threefold symmetry axis marked by a triangle. **g** RMSD table of the mutated HA ectodomain relative to the wild-type. RMSD values were calculated for five residues neighboring each mutation site to compare the wild-type and mutant HA proteins.

We measured the thermal stability of the mutated protein using the DSF method (Fig. 9c). The mutation gave a 4.9-degree increase of Tm values at pH 8.0 and 4.7 degrees at pH 5.5, indicating successful stabilization of the protein structure by the mutation. The structure of the mutated H9/HK98 HA proteins was evaluated using cryo-EM, and the structure was refined at 2.9 Å resolution. As shown in Fig. 9d, the 2D class-averaged maps clearly illustrate the trimeric pre-fusion structure. The refined structure was superimposable to that of the previously reported pre-fusion structure with PDB id 1JSD (Fig. 9e). A close-up view near the mutated sites indicates that the overall structure of the protein was not disturbed by the mutations (Fig. 9f and g).

### Stabilization of trimeric states of other group 1 influenza HA proteins

To evaluate whether our five-site optimization strategy can stabilize HA trimers from other group 1 influenza viruses, specifically those beyond H1 and H9, we redesigned six HA proteins from H5, H6, H8, H12, H13, and H18 (Supplementary Figs. 6-11). Ectodomains of the HA proteins from A/Vietnam/1194/2004 (H5N1), A/Taiwan/2/2013 (H6N1), /turkey/Ontario/6118/1968 (H8N4), A/duck/Alberta/60/1976 (H12N5), A/gull/Maryland/704/1977 (H13N6), A/flat-faced bat/Peru/033/2010 (H18N11) viruses were used for the experiments. The five amino acid residues selected in the H1/Cal09 study in each HA protein were analyzed using the protein MPNN program, and the optimal mutations were selected for these sites (Fig. 10a). The previously reported structures with PDB id, 4BGW, 5BR0, 6V46, 7A9D, 4KPQ and 4K3X were used as the structural templates for the protein MPNN calculation ^1, 23–27^. We then produced both the wild-type and mutated HA ectodomains and analyzed their trimerization states using gel permeation chromatography at pH 8.0. As shown in Fig. 10b, the wild-type ectodomains of H5 and H18 HA proteins primarily formed monomers in solution. In contrast, the mutated proteins were eluted as trimers, demonstrating that the mutagenesis effectively stabilized the trimeric structure. For H8 and H13, the wild-type ectodomains exhibited a mixture of monomers and trimers in solution. After mutation, the proteins were nearly entirely trimeric, and monomer peaks were not detectable. Regarding H6 and H12 HA proteins, the wild-type ectodomains could not be produced in the insect cells, so only the mutated proteins, which were mainly eluted as trimers, were analyzed. This suggests not only that the mutations stabilized the trimeric states but also that the mutations enhanced protein expression in the insect cells. In conclusion, we have shown that the five-site optimization strategy reliably stabilizes the trimeric pre-fusion structures of most of the group 1 HA proteins.

**Fig. 10.**
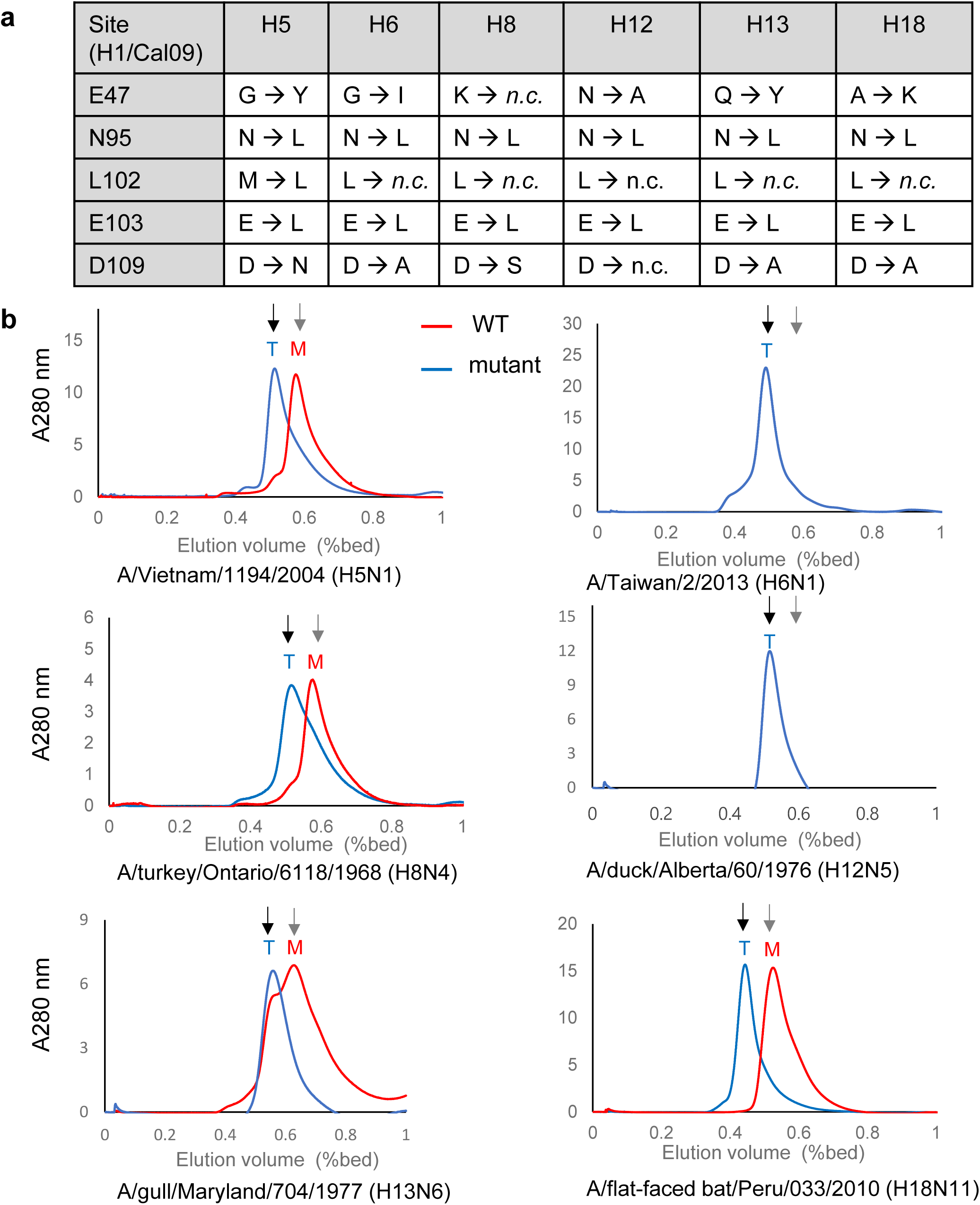
Trimerization states of the mutated group 1 HA ectodomains. **a** Summary of the mutations introduced into the ectodomains of group 1 HA proteins. *n.c.*, no change. **b** Gel permeation elution profiles of the ectodomains from group 1 influenza HA proteins. The expected elution positions of the trimeric and monomeric forms are indicated with ‘T’ for trimer and ‘M’ for monomer, respectively. Elution volumes of β-amylase (200 kDa) and BSA (66 kDa) are indicated by black and gray arrows, respectively.

## Discussion

In this study, we demonstrated that computational optimization of a small number of amino acid residues in the trimerization interface can effectively stabilize the pre-fusion structure of most of the group 1 influenza HA proteins. By targeting five suboptimal amino acid residues in the trimerization interface of the HA proteins, we could engineer mutant HA proteins from eight subgroups of influenza viruses that showed improved trimerization and protein stability. High-resolution cryo-EM studies confirmed their adoption of stable pre-fusion structures, underscoring the potential of our strategy for the rapid stabilization of HA pre-fusion structures. This approach is particularly pertinent in the context of influenza’s antigenic drift and shift, which complicates vaccine development and necessitates effective strategies to enhance cross-reactive immunity.

Our approach diverges from that of Milder *et al.*, who achieved stabilization through the H26W, K51I and E103I mutations at pH-sensitive switch sites of H1 HA ^28^. Their mutational sites are different from ours and their methodology did not use computational redesign of the amino acid residues. Nevertheless, their mutations could increase the thermal stability of the engineered HA protein by more than ∼10 degrees and be applied to a broad spectrum of HA proteins. We believe that having multiple sets of stabilizing mutations is crucial for pandemic preparedness, especially against influenza and other viruses with unpredictable sequence variations. Both Milder *et al.*’s and our selection methods of mutational sites need improvement. It is based on manual structural inspection, a subjective process that requires more objective and computationally driven criteria in future research.

Traditional studies on HA pre-fusion structures primarily rely on X-ray crystallography, which demands the crystallization of protein samples at high protein concentrations in non-physiological conditions that often contain high concentrations of precipitating reagents like ammonium sulfate or polyethylene glycols ^29^. This can lead to misleading conclusions, as such conditions may induce the trimerization of otherwise monomeric proteins at more physiologically relevant protein concentrations and buffer conditions. Low-resolution techniques like negatively stained EM also have limitations in confirming the stabilization of pre-fusion structures because they cannot detect small but significant structural deviations in the proteins, as seen in Fig. 2c. Our work highlights the usefulness of high-resolution cryo-EM for definitive structural confirmation, advocating for its broader application in future studies.

The potential threat of an influenza pandemic, particularly from avian influenza viruses in H5N1, underscores the urgency of preparedness efforts ^30, 31^. These viruses’ ability to mutate and reassort emphasizes their pandemic potential. Our research suggests that once a set of suboptimal residues is experimentally identified in advance, rapid redesign of HA sequences from novel viral strains, even with significant sequential variation, is feasible using computational tools like the protein MPNN program. The computational calculation can be achievable within hours with moderate computational resources. Therefore, our method can be a strategic response to emerging influenza strains with unexpected sequential variations. Continuous efforts to identify multiple sets of key mutational sites will further enhance readiness for future outbreaks of influenza and other zoonotic viruses.

## Methods

### Sequence optimization using Protein MPNN

The previously reported structures of pre-fusion forms of HA from human group 1 influenza were used as structural input to Protein MPNN, and five amino acid positions selected at the trimerization interface were redesigned (Supplementary Figs. 1 and 2). For protein MPNN calculation, a temperature equal to 0.1 and the random seed were used to generate outputs. 10,000 sequences were generated with the imposition of a trimeric symmetry. The sequences calculated with the lowest scores were selected for structure prediction and manual inspection. The structures were predicted with AlphaFold2 with 5 recycling steps. The best models predicted by AlphaFold2 were manually evaluated and selected for experimental testing.

### Expression and purification of H1/Cal09 and H9/HK98 HA

The genes encoding the ectodomains of H1/Cal09 and H9/HK98 HAs were synthesized (Twist Bioscience) and cloned into a pEG BacMam baculovirus transfer vector (Supplementary Tables 1 and 2) ^32^. A PreScission protease site, mCherry, thrombin cleavage site, and ALFA tag were added to the C-terminus of HAs. The plasmid was transformed into DH10Bac *E.coli* (ThermoFisher) for transposition into a bacmid. The recombinant baculovirus was generated by transfecting Sf9 insect cells with the bacmid DNA. Protein was expressed in Expi293F cells cultured in the Expi293 media (ThermoFisher) in an 8% CO_2_ shaking incubator. The cells were infected by treating 10% (v/v) baculovirus at a density of 3×10^6^ cells/ml. Protein expression was enhanced by treating with 10 mM sodium butyrate (Sigma-Aldrich) after 18∼22 hours of infection and further incubated for 5 to 7 days. The supernatant was collected by centrifugation at 4,500 rpm for 20 minutes and loaded onto a column packed with an agarose resin conjugated to an anti-ALFA nanobody ^33^. The anti-ALFA-nanobody column was washed with 20 column volumes of wash buffer containing 20 mM Tris-HCl pH 8.0, 200 mM NaCl. Protein was eluted by treating with 3% (w/w) thrombin (Lee Biosolutions) overnight. The eluted protein solution was concentrated using an ultracentrifugal filter with the 30 kDa cutoff (Merck Milipore). The protein was further purified by a Superdex 200 increase 10/300 GL gel filtration column (Cytiva) equilibrated with a buffer containing 20 mM Tris-HCl pH 8.0, 200 mM NaCl.

### Expression and purification of H1/Mal07 and other group 1 HAs

The genes encoding the ectodomains of HAs for H5 and H9 were synthesized by Gene Universal, while other subtypes were synthesized by Twist Bioscience, and all were cloned into a pAcGP67a baculovirus transfer vector (ThermoFisher) (Supplementary Tables 3-7). Recombinant baculovirus was generated by co-transfection with a linearized baculovirus genome, BestBac2.0 (Expression Systems), into Sf9 insect cells. Protein was expressed in High Five insect cells cultured in ESF 921 media (Expression Systems) by adding 3∼4% (v/v) baculovirus at 21℃ for 72 hours. The secreted protein was bound to cOmplete His-Tag Purification column (Roche) and eluted using an elution buffer containing 20 mM Tris-HCl, 200 mM NaCl, and a gradient of 100 mM to 500 mM imidazole at pH 8.0. The protein was further purified by a Superdex 200 increase 10/300 GL gel filtration column (Cytiva) equilibrated with a buffer containing 20 mM Tris-HCl pH 8.0, 200 mM NaCl.

### Mouse immunization

Six-week-old female BALB/c mice were purchased from Daehan Biolink (Eumseong, Korea). Five mice per group were immunized three times via intramuscular (IM) injection in the outer thigh. For the initial immunization, 5 ug of antigen was mixed 1:1 with Freund’s complete adjuvant. Two weeks later, a second dose, 5ug of antigen mixed 1:1 with Fruend’s incomplete adjuvant, was injected. One week after the second injection, a final dose of 5 ug antigen was injected without adjuvant. Serum samples were collected by retro-orbital bleeding one week after the final immunization. All animal procedures were approved and guided by the Institutional Animal Care and Use Committee (IACUC) of Pohang University of Science and Technology (POSTECH-2024-0066).

### Enzyme-linked immunosorbent assay (ELISA)

Serum levels of HA-specific antibodies were determined by ELISA. Wells of 96-well microtiter plates (Komabiotech., Seoul, Korea) were coated with 90 ul of 5 ug/ml recombinant HA in coating buffer, pH 9.6 (Komabiotech) at 4℃ overnight. Plates were washed 3 times with washing buffer, 1X PBS including 0.05 % Tween 20, and blocked with 2% skim milk in PBS for 2 h at RT. Mouse sera were diluted 1:100 in PBS with 0.1% BSA, and then serial 10-fold dilutions were prepared and added to the wells in triplicate. After incubation for 2h at RT on a shaker, plates were washed 3 times with washing buffer, and horseradish peroxide (HRP)- conjugated anti-mouse IgG (Abcam, Cambridge, UK) diluted 1:10,000 in PBS with 0.1% BSA was added. Following 2h incubation at RT, the plates were washed and 100 ul TMB developing solution (Komabiotech) was added to each well. The Reaction was stopped after 5 min by adding 100 ul of 0.5 M sulfuric acid. Absorbance was measured at 450 nm using a microplate reader (Tecan GENios Pro, Mannedorf, Switzerland). The dilutions of the mouse sera that gave two-fold higher ELISA signals than the sera of pre-immunized mice at 1:100 dilution were designated the endpoint titers.

### Preparation of cryo-EM grids

Purified HAs at 0.6 ∼ 1.0 mg/ml concentration were prepared. The 300-mesh UltraAuFoil R 1.2/1.3 EM grids (Structure Probe) were discharged at 15 mA for 60 seconds using a glow-discharger (PELCO). All EM grids were prepared by applying 3.2 μl of protein sample to the EM grids, followed by 4 seconds of blotting and 30 seconds of waiting time using the Vitrobot Mark IV (ThermoFisher) under 4°C and 100% humidity, before plunge freezing into liquid ethane.

### Cryo-EM data collection

The data collection statistics are summarized in Supplementary Tables S8 and S9. The H1/Cal09-WT, H1/Cal09-WT(tri), H1/Mal07-WT and H1/Mal07-WT(tri) data sets were collected using a Titan Krios microscope at 300 kV, equipped with an energy filter and a Falcon4i direct electron detector (ThermoFisher) operating in counting mode at 130,000x magnification with a pixel size of 0.938 Å. The H1/Cal09-mut, H1/Mal07-5mut and H9/HK98-mut HA data sets were collected using a Titan Krios microscope (ThermoFisher) at 300 kV, equipped with an energy filter and a K3 direct electron detector (Gatan) operating in counting mode. H1/Cal09-mut data were acquired at 105,000x magnification operating with a pixel size of 0.85 Å, while other data sets were collected at 135,000x magnification with a pixel size of 0.651 Å. Each cryo-EM movie was recorded in the TIFF format with a total dose of 70 e/Å². All data sets were automatically acquired using EPU software for single particle analysis (ThermoFisher)

### Cryo-EM 2D data processing

All cryo-EM data processing was performed using the CryoSPARC v4.6.0 program ^34^ and GPGPU cluster at the Institute of Membrane Proteins (NFEC-2025-03-304437). The cryo-EM movie files were preprocessed using the patch motion correction and the patch contrast transfer function (CTF) methods. The preprocessed data were imported into the CryoSPARC program with an up-sampling factor of 1, followed by patch motion correction and patch CTF estimation. Poor-quality micrographs were manually curated based on the CTF estimations, ice thickness, and the total motion of the frames. Initial particles were picked using the Topaz model and purified by the 2D class average method. The high-quality 2D class averages were selected for Topaz picking model generation ^35^. The particle picking method is further optimized by repeating 2∼3 rounds of Topaz analysis and multi-rounds of 2D classification.

Data processing for trimeric H1/Cal09-mut, particles were extracted with a box size of 320 pixels and binned by 100 pixels (Supplementary Fig. 12). The initial model was generated using *Ab-initio* reconstruction with C1 symmetry, followed by particle re-centering and un-binning. The electron density map was refined through non-uniform refinements with C3 symmetry. The noise in the final map was reduced using DeepEMhancer, an automatic deep learning-based sharpening method ^36^.

Data processing for trimeric H1/Mal07-mut, particles were extracted with a box size of 420 pixels and binned by 100 pixels (Supplementary Fig. 13). The initial model was generated using *Ab-initio* reconstruction with C1 symmetry, followed by particle re-centering and un-binning. To obtain a more refined electron density map, additional particle extraction was performed using Topaz training and Topaz extract, and the map was subsequently refined using non-uniform refinement with C3 symmetry. The noise in the final map was reduced using the DeepEMhancer program.

Data processing for trimeric H9/HK98 HA mutant, particles were extracted with a box size of 420 pixels and binned by 100 pixels (Supplementary Fig. 14). The initial model was generated using *Ab-initio* reconstruction with C1 symmetry, followed by particle re-centering and un-binning. To obtain a more refined electron density map, additional particle extraction was performed using Topaz training and Topaz extract, and the map was subsequently refined using non-uniform refinement with C3 symmetry. The noise in the final map was reduced using the DeepEMhancer program.

### Model building

The initial model for H1/Cal09-mut was generated by AlphaFold2 ^37^, while the models for H1/Mal07-mut and H9/HK98-mut were generated by AlphaFold3 ^38^. These models were fitted into the cryo-EM density map using the ChimeraX program ^39^. The resulting structure was refined through multiple rounds of manual model building using the Coot program and real-space refinement using the Phenix program ^40, 41^.

### Differential scanning fluorometry (DSF) and gel permeation chromatography

For the H1/Cal09 and H1/Mal07 HA proteins, denaturation temperatures, Tm, were determined using a Prometheus NT.48 instrument (NanoTemper). Samples were prepared at a concentration of 0.5 mg/mL in a buffer containing 20 mM Tris-HCl pH 8.0, 200 mM NaCl and loaded into DSF capillaries and subjected to thermal stress from 25 °C to 95 °C at a heating rate of 1 °C/min. Fluorescence emission from tryptophan residues following UV excitation at 280 nm was collected at 330 nm and 350 nm using a dual-UV detector. Thermal stability parameters, including T*onset* and Tm, were calculated using PR.ThermControl software (NanoTemper).

For the H9/HK98 HA protein, DSF analysis using intrinsic tryptophan fluorescence with the Prometheus instrument yielded a large negative peak, which complicated the accurate measurement of melting temperatures. Therefore, the analysis method was changed to DSF using Protein Thermal Shift dye (Thermo Fisher). In this method, the fluorescence signal from the dye increases upon exposure of hydrophobic amino acid residues during protein denaturation. For analysis, samples were prepared at a concentration of 0.3 mg/mL in a buffer containing 20 mM Tris and 200 mM NaCl, pH 8. Fluorescence signals with an excitation wavelength of 580 nm and an emission wavelength of 623 nm were measured using a QuantStudio 3 real-time PCR system (Thermo Fisher) across a temperature range from 25 °C to 95 °C at a heating rate of 0.05 °C/min.

For gel permeation chromatography, purified HA proteins were diluted to 0.2 mg/mL in buffers containing either 20 mM sodium acetate, pH 4.7 or 5.5 with 75 mM NaCl, or 20 mM Tris-HCl, pH 8.0 with 200 mM NaCl. The protein samples were incubated for 30 minutes at 4 °C and subsequently loaded onto Superdex 200 Increase 10/300 or Superdex 200 Increase 5/150 columns.

## Data availability

The cryo-EM maps and the atomic coordinates will be deposited in the Electron Microscopy Data Bank (https://www.ebi.ac.uk/pdbe/emdb) and the Protein Data Bank (https://www.rcsb.org/), respectively, under the following accession codes: EMD-64329 and 9UMY for the H1/Cal09-mut HA, EMD-64334 and 9UN0 for the H1/Mal07-mut HA; EMD-64336 and 9UN1 for the H9/HK98-mut HA.

## Code availability

This paper does not report the original code.

## Availability of materials

All unique materials used are readily available from the authors.

## Author Contributions

J.-O.L. conceived and supervised the project. G.Y.L. and Y.G.K. performed computational optimization of HA sequences, expressed and purified proteins, prepared samples for cryo-EM studies, collected and processed cryo-EM data, and built and refined atomic models. C.J.L. improved the cryo-EM map using computational methods. G.Y.L., Y.G.K., and J.-O.L. wrote and edited the manuscript with input from all authors.

