## Supplementary material for "Stabilization of the trimeric pre-fusion structures of influenza H1 and H9 hemagglutinins by mutations in the stem helices": Supplemenatary Figures and Tables

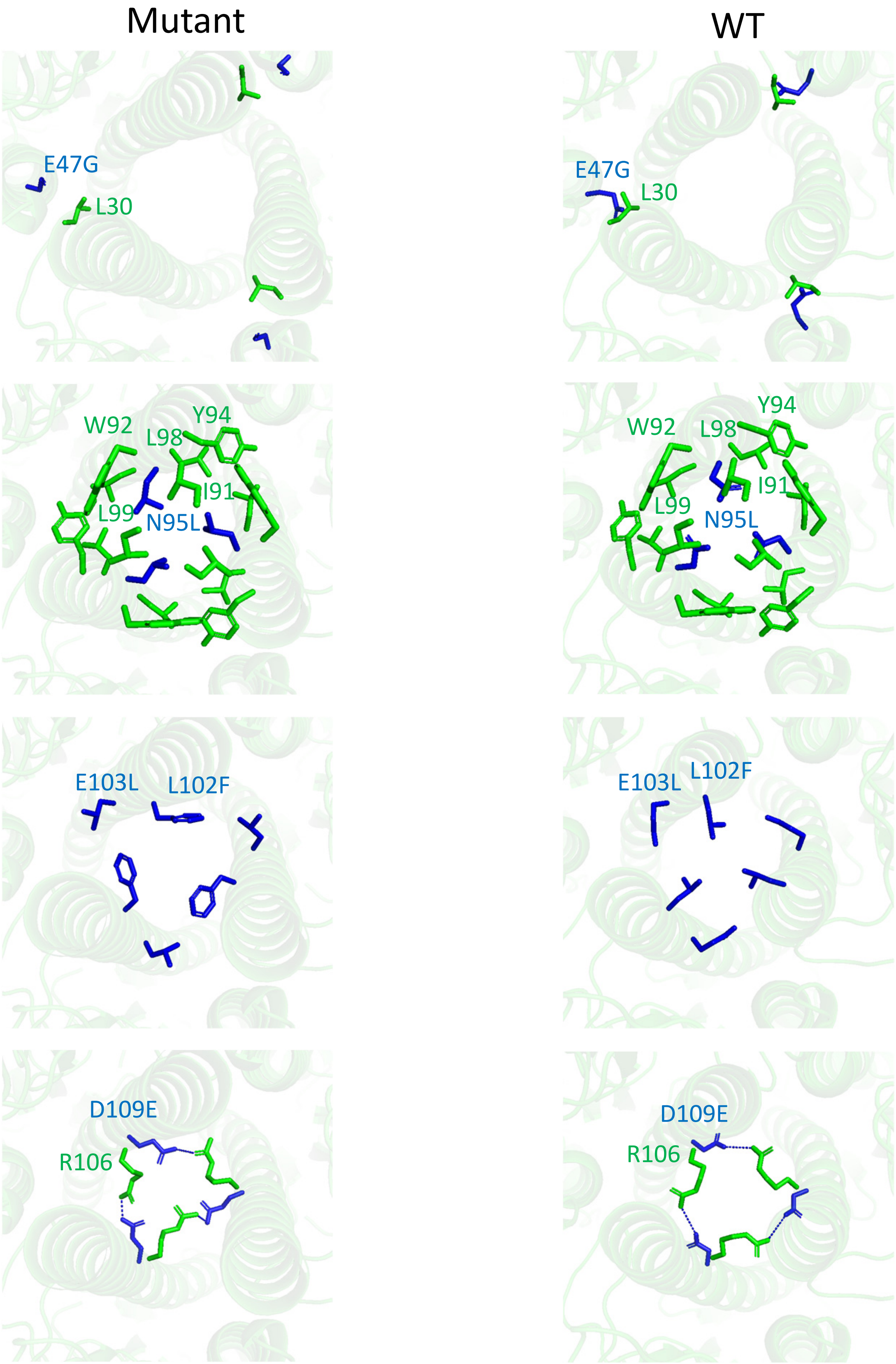

**Supplementary Fig. x | Close-up view of the mutated sites for H1/Cal09-mut**

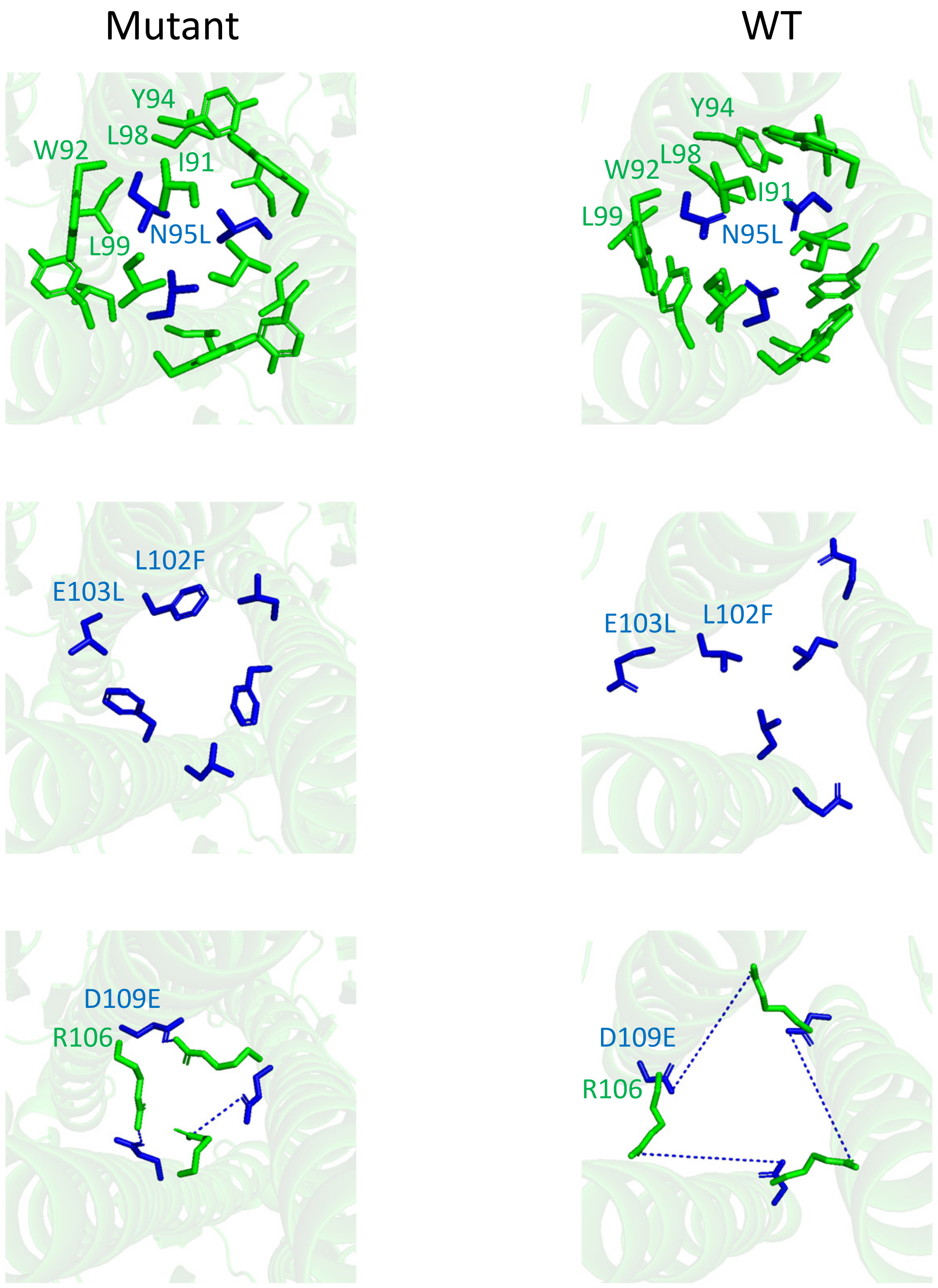

**Supplementary Fig. x | Close-up view of the mutated sites for H1/Mal07-mut**

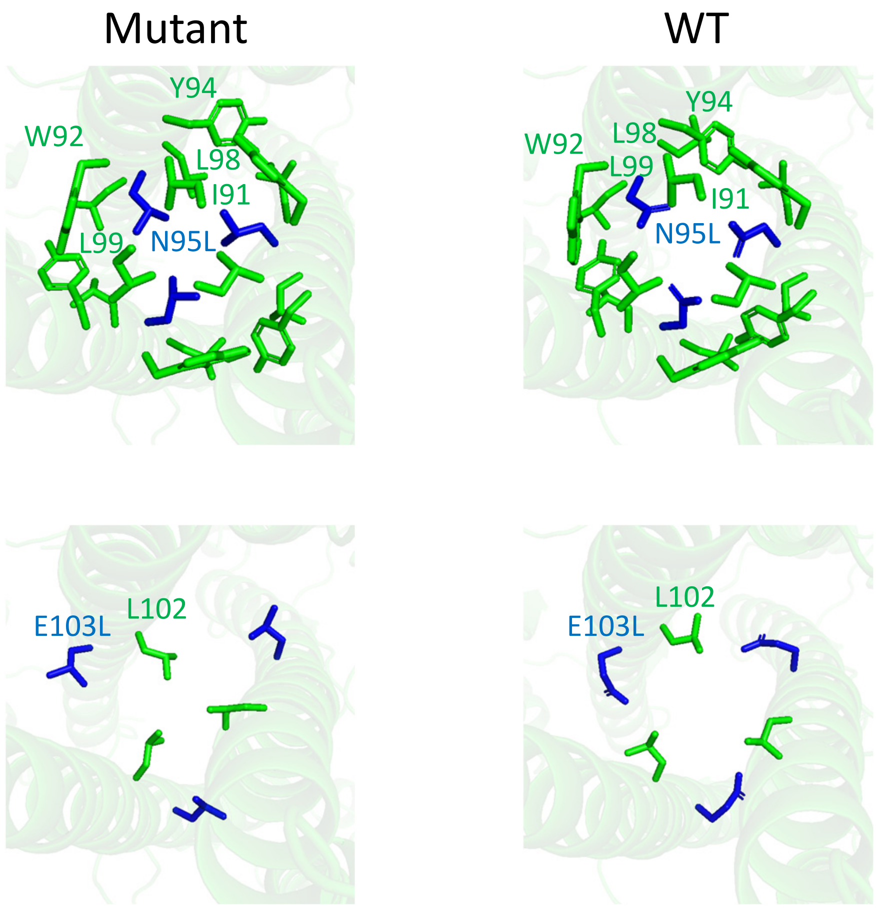

**Supplementary Fig. x | Close-up view of the mutated sites for H9/HK98-mut**

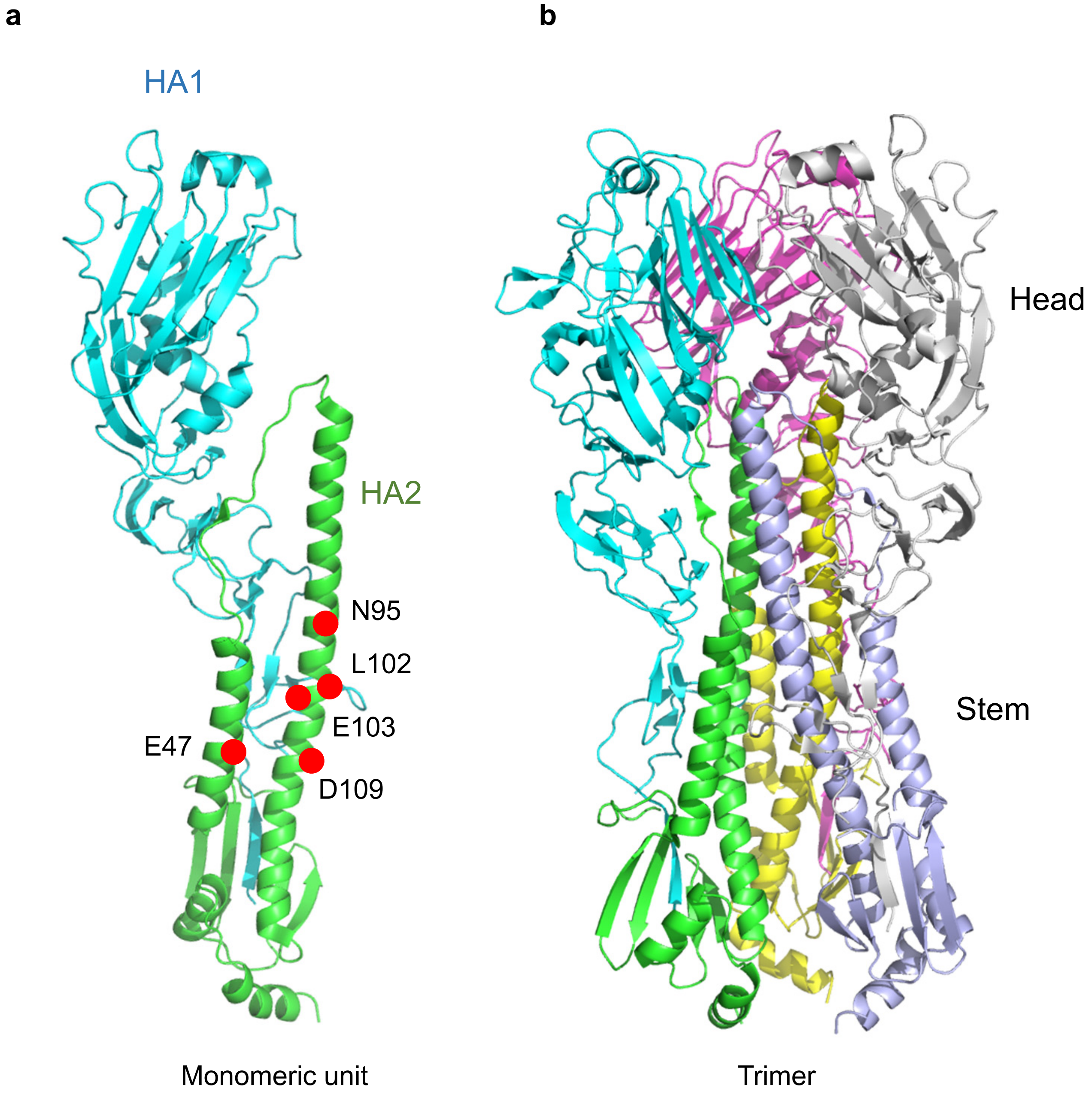

**Supplementary Fig. 1 | Positions of the computationally optimized amino acid sites. a** Structure of a single monomer within the HA trimer, shown in the left panel. Mutated amino acid sites are indicated with red dots. **b** Structure of the full HA trimer. The view is rotated approximately 30 degrees along the vertical axis relative to panel **a**.

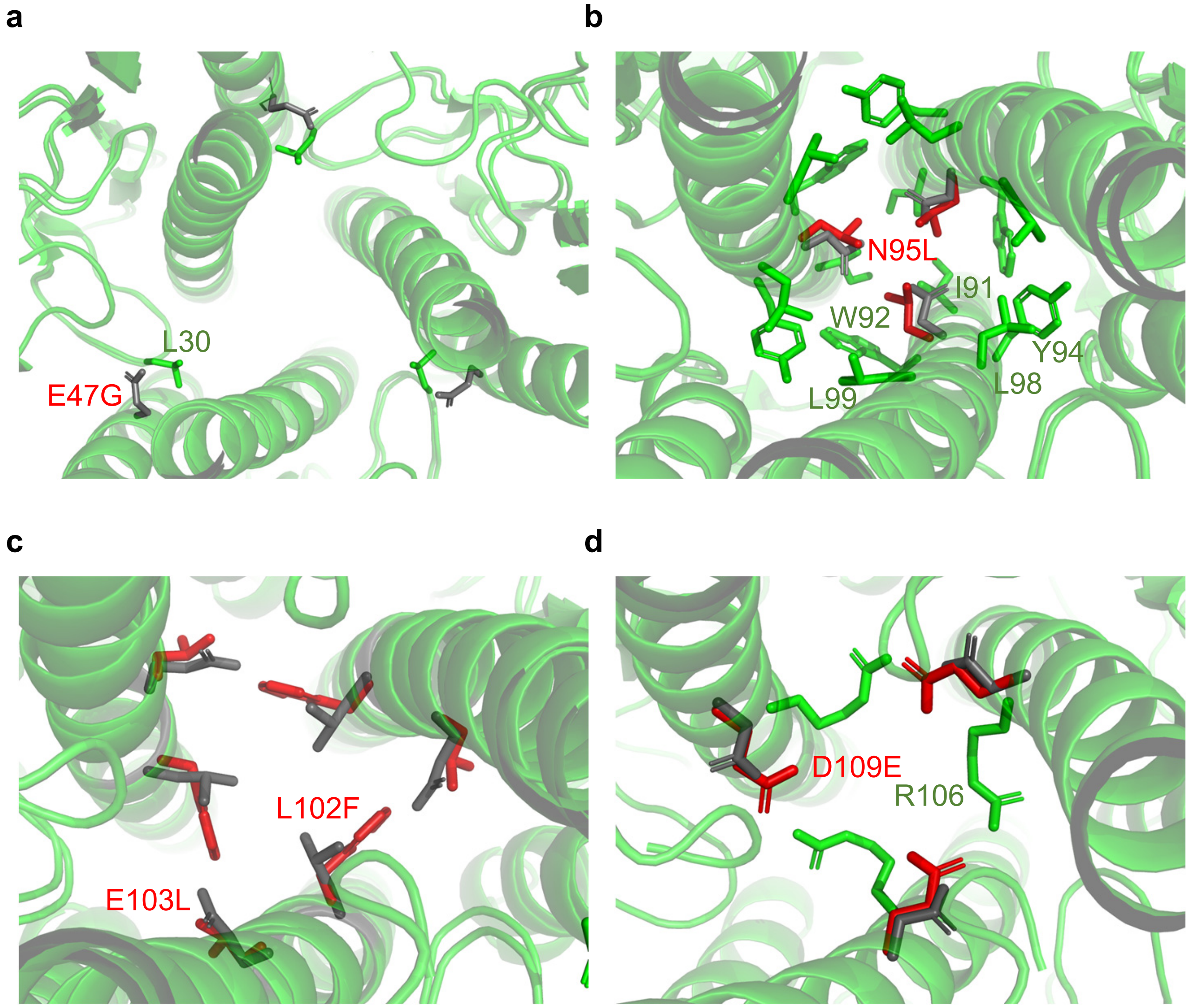

**Supplementary Fig. 2 | Close-up view of the computationally optimized sites.** The view is from above, looking down the central stem helices. Mutated residues are labeled in red. Side chains from the wild-type structure are shown in dark grey, and those from the mutated structure are shown in red. Amino acid residues that interact with the mutated sites are shown in green. One of the three HA subunits in the HA trimer is labeled.

```
H1/Mal07        DTICIGYHANNSTDTVDTVLEKNVTVTHSVNLLEDSHNGKLCLLKGIAPLQLGNCSVAGW
H1/Cal09        DTLCIGYHANNSTDTVDTVLEKNVTVTHSVNLLEDKHNGKLCKLRGVAPLHLGKCNIAGW
                **:********************************.****** *:*:***:**:*.:***

H1/Mal07        ILGNPECELLISRESWSYIVEKPNPENGTCYPGHFADYEELREQLSSVSSFERFEIFPKE
H1/Cal09        ILGNPECESLSTASSWSYIVETPSSDNGTCYPGDFIDYEELREQLSSVSSFERFEIFPKT
                ******** * : .*******.*..:*******.* ***********************

H1/Mao07        SSWPNHTTTG-VSASCSHNGESSFYKNLLWLTGKNGLYPNLSKSYANNKEKEVLVLWGVH
H1/Cal09        SSWPNHDSNKGVTAACPHAGAKSFYKNLIWLVKKGNSYPKLSKSYINDKGKEVLVLWGIH
                ****** :.  *:*:*.* * .******:**. *.. **:***** *:* ********:*

H1/Mal07        HPPNIGDQRALYHTENAYVSVVSSHYSRKFTPEIAKRPKVRDQEGRINYYWTLLEPGDTI
H1/Cal09        HPSTSADQQSLYQNADTYVFVGSSRYSKKFKPEIAIRPKVRDQEGRMNYYWTLVEPGDKI
                **.. .**::**:. ::** * **:**:**.**** **********:******:****.*

H1/Mal07        IFEANGNLIAPRYAFALSRGFGSGIINSNAPMDECDAKCQTPQGAINSSLPFQNVHPVTI
H1/Cal09        TFEATGNLVVPRYAFAMERNAGSGIIISDTPVHDCNTTCQTPKGAINTSLPFQNIHPITI
                 ***.***:.******:.*. ***** *::*:.:*::.****:****:******:**:**

H1/Mal07        GECPKYVRSAKLRMVTGLRNIPSIQSRGLFGAIAGFIEGGWTGMVDGWYGYHHQNEQGSG
H1/Cal09        GKCPKYVKSTKLRLATGLRNIPSIQSRGLFGAIAGFIEGGWTGMVDGWYGYHHQNEQGSG
                *:*****:*:***:.*********************************************

H1/Mal07        YAADQKSTQNAINGITNKVNSVIEKMNTQFTAVGKEFNKLERRMENLNKKVDDGFIDIWT
H1/Cal09        YAADLKSTQNAIDEITNKVNSVIEKMNTQFTAVGKEFNHLEKRIENLNKKVDDGFLDIWT
                **** *******: ************************:**:*:***********:****

H1/Mal07        YNAELLVLLENERTLDFHDSNVKNLYEKVKSQLKNNAKEIGNGCFEFYHKCNDECMESVK
H1/Cal09        YNAELLVLLENERTLDYHDSNVKNLYEKVRSQLKNNAKEIGNGCFEFYHKCDNTCMESVK
                ****************:************:*********************:: ******

H1/Mal07        NGTYDYPKYSEESKLNREKID
H1/Cal09        NGTYDYPKYSEEAKLNREEID
                ************:*****:**
```

**Supplementary Fig. 3 | Sequence alignment of HA ectodomains from H1/Mal07 and H1/Cal09.** The boundary of the HA1 and HA2 chains is marked with an arrow. The amino acid sites that are computationally optimized are highlighted with red boxes.

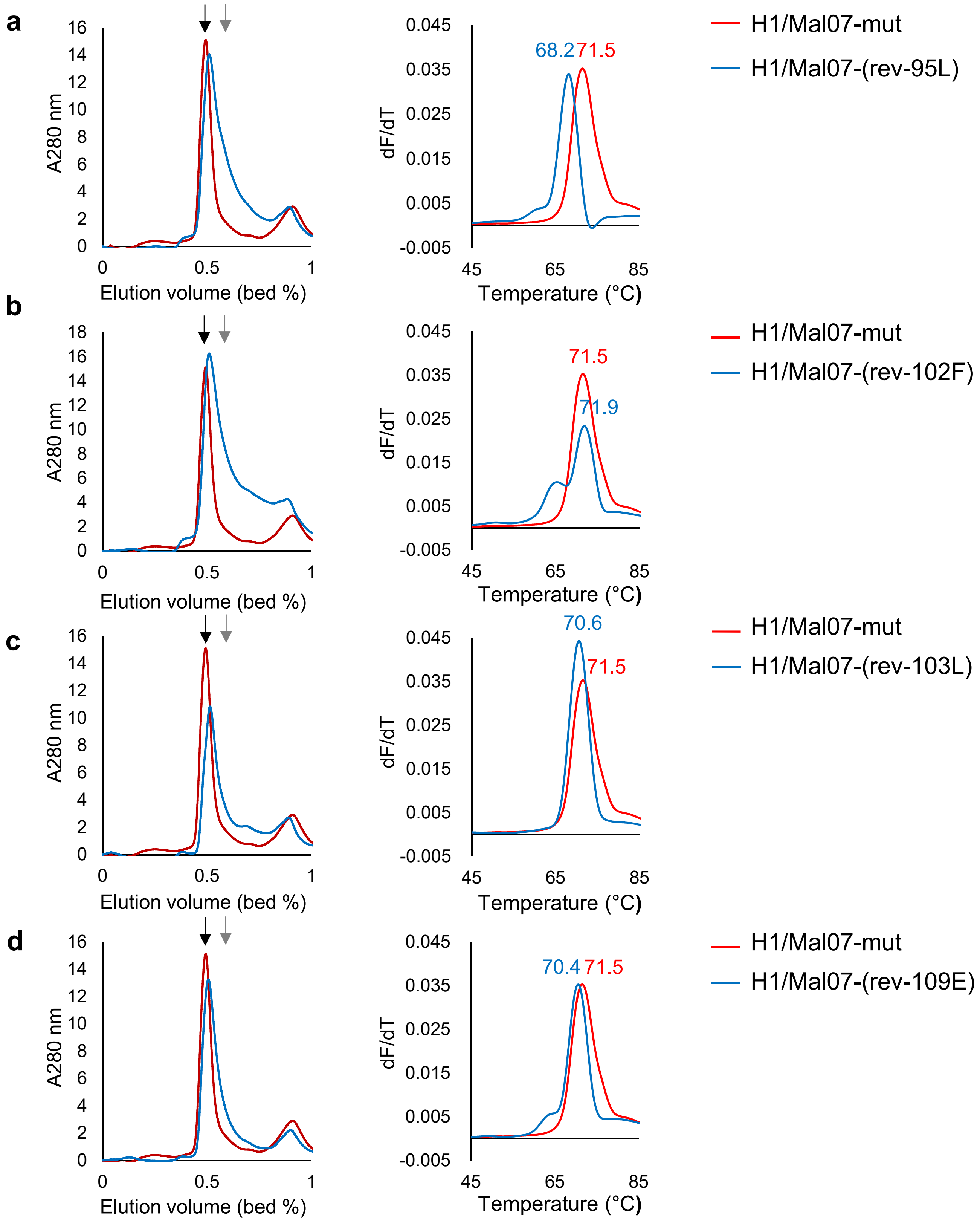

**Supplementary Fig. 4 | Trimerization state and thermal stability at pH 5.5 of wild-type and mutant H1/Mal07 HA ectodomains. a-d** Gel permeation chromatography (left) and differential scanning fluorimetry (DSF) profiles (right) of H1/Mal07 HA containing all four mutations (H1/Mal07-mut) or containing only three of the four mutations, with one mutation reverted back to the wild-type sequence. All experiments were conducted at pH 5.5. Elution volumes of β-amylase (200 kDa) and BSA (66 kDa) are indicated by black and gray arrows, respectively.

```
H9/HK98         DKICIGYQSTNSTETVDTLTETNVPVTHAKELLHTSHNGMLCATNLGHPLILDTCTIEGL
H1/Cal09        DTLCIGYHANNSTDTVDTVLEKNVTVTHSVNLLEDKHNGKLCKLRGVAPLHLGKCNIAGW
                *.:****::.***:****: *.**.***: :**. .*** **  .   ** *..*.* *

H9/HK98         IYGNPSCDLLLGGREWSYIVERPSAVNGMCYPGNVENLEELRSLFSSASSYQRIQIFPDT
H1/Cal09        ILGNPECESLSTASSWSYIVETPSSDNGTCYPGDFIDYEELREQLSSVSSFERFEIFPKT
                * ***.*: *  . .****** **: ** ****:. : ****. :**.**::*::***.*

H9/HK98         IWNVSYSG-------TSSACSDSFYRSMRWLTQKNNAYPIQDAQYTNNRGKSILFMWGIN
H1/Cal09        SSWPNHDSNKGVTAACPHAGAKSFYKNLIWLVKKGNSYPKLSKSYINDKGKEVLVLWGIH
                    .:..        . * :.***:.: **.:*.*:**  . .* *::**.:*.:***:

H9/HK98         HPPTDTVQTNLYTRTDTTTSVTTEDINRTFKPVIGPRPLVNGLHGRIDYYWSVLKPGQTL
H1/Cal09        HPSTSADQQSLYQNADTYVFVGSSRYSKKFKPEIAIRPKVRDQEGRMNYYWTLVEPGDKI
                **.*.: * .** .:** . * :.  .:.*** *. ** *.. .**::***::::**:.:

H9/HK98         RVRSNGNLIAPWYGHILSGESHGRILKTDLNSGNCVVQCQTERGGLNTTLPFHNVSKYAF
H1/Cal09        TFEATGNLVVPRYAFAMERNAGSGIIISDTPVHDCNTTCQTPKGAINTSLPFQNIHPITI
                 ..:.***:.* *.. :. :: . *: :*    :* . *** :*.:**:***:*:   ::

H9/HK98         GNCPKYVGVKSLKLAVGLRNVPARSSRGLFGAIAGFIEGGWPGLVAGWYGFQHSNDQGVG
H1/Cal09        GKCPKYVKSTKLRLATGLRNIPSIQSRGLFGAIAGFIEGGWTGMVDGWYGYHHQNEQGSG
                *:*****  ..*:**.****:*: .* **************.*:* ****::*.*:** *

H9/HK98         MAADSDSTQKAIDKITSKVNNIVDKMNKQYGIIDHEFSEIETRLNMINNKIDDQIQDIWT
H1/Cal09        YAADLKSTQNAIDEITNKVNSVIEKMNTQFTAVGKEFNHLEKRIENLNKKVDDGFLDIWT
                 *** .***:***:**.***.:::***.*:  :.:**..:*.*:: :*:*:** : ****

H9/HK98         YNAELLVLLENQKTLDEHDANVNNLYNKVKRALGSNAMEDGKGCFELYHKCDDQCMETIR
H1/Cal09        YNAELLVLLENERTLDYHDSNVKNLYEKVRSQLKNNAKEIGNGCFEFYHKCDNTCMESVK
                ***********::*** **:**:***:**:  * .** * *:****:*****: ***:::

H9/HK98         NGTYNRRKYKEES
H1/Cal09        NGTYDYPKYSEEA
                ****:  **.**:
```

**Supplementary Fig. 5 | Sequence alignment of HA ectodomains from H9/HK98 and H1/Cal09.** The boundary of the HA1 and HA2 chains is marked with an arrow. The amino acid sites that are computationally optimized are highlighted with red boxes.

```
H1/Cal09        DTLCIGYHANNSTDTVDTVLEKNVTVTHSVNLLEDKHNGKLCKLRGVAPLHLGKCNIAGW
H5              DQICIGYHANNSTEQVDTIMEKNVTVTHAQDILEKTHNGKLCDLDGVKPLILRDCSVAGW
                * :**********: ***::********: ::**..******.* ** ** * .*.:***

H1/Cal09        ILGNPECESLSTASSWSYIVETPSSDNGTCYPGDFIDYEELREQLSSVSSFERFEIFPKT
H5              LLGNPMCDEFINVPEWSYIVEKANPVNDLCYPGDFNDYEELKHLLSRINHFEKIQIIPKS
                :**** *:.: ....******.... *. ****** *****:. ** :. **:::*:**:

H1/Cal09        SSWPNHDSNKGVTAACPHAGAKSFYKNLIWLVKKGNSYPKLSKSYINDKGKEVLVLWGIH
H5              S-WSSHEASLGVSSACPYQGKSSFFRNVVWLIKKNSTYPTIKRSYNNTNQEDLLVLWGIH
                * *..*::. **::***: * .**::*::**:**..:**.:.:** * : :::*******

H1/Cal09        HPSTSADQQSLYQNADTYVFVGSSRYSKKFKPEIAIRPKVRDQEGRMNYYWTLVEPGDKI
H5              HPNDAAEQTKLYQNPTTYISVGTSTLNQRLVPRIATRSKVNGQSGRMEFFWTILKPNDAI
                **. :*:* .****. **: **:*  .::: *.** *.**..*.***:::**:::*.* *

H1/Cal09        TFEATGNLVVPRYAFAMERNAGSGIIISDTPVHDCNTTCQTPKGAINTSLPFQNIHPITI
H5              NFESNGNFIAPEYAYKIVKKGDSTIMKSELEYGNCNTKCQTPMGAINSSMPFHNIHPLTI
                .**:.**::.*.**: : ::..* *: *:    :***.**** ****:*:**:****:**

H1/Cal09        GKCPKYVKSTKLRLATGLRNIPSIQSRGLFGAIAGFIEGGWTGMVDGWYGYHHQNEQGSG
H5              GECPKYVKSNRLVLATGLRNSPQRETRGLFGAIAGFIEGGWQGMVDGWYGYHHSNEQGSG
                *:*******.:* ******* *. ::*************** ***********.******

H1/Cal09        YAADLKSTQNAIDEITNKVNSVIEKMNTQFTAVGKEFNHLEKRIENLNKKVDDGFLDIWT
H5              YAADKESTQKAIDGVTNKVNSIIDKMNTQFEAVGREFNNLERRIENLNKKMEDGFLDVWT
                **** :***:*** :******:*:****** ***:***:**:********::*****:**

H1/Cal09        YNAELLVLLENERTLDYHDSNVKNLYEKVRSQLKNNAKEIGNGCFEFYHKCDNTCMESVK
H5              YNAELLVLMENERTLDFHDSNVKNLYDKVRLQLRDNAKELGNGCFEFYHKCDNECMESVR
                ********:*******:*********:*** **::****:************* *****:

H1/Cal09        NGTYDYPKYSEEA
H5              NGTYDYPQYSEEA
                *******:*****
```

**Supplementary Fig. 6 | Sequence alignment of HA ectodomains from H5 and H1/Cal09.** The boundary of the HA1 and HA2 chains is marked with an arrow. The amino acid sites that are computationally optimized are highlighted with red boxes.

```
H1/Cal09        DTLCIGYHANNSTDTVDTVLEKNVTVTHSVNLLEDKHNGKLCKLRGVAPLHLGKCNIAGW
H6              DKICIGYHANNSTTQVDTLLEKNVTVTHSVELLENQKEKRFCKIMNKAPLDLKDCTIEGW
                *.:**********  ***:***********:***:::: ::**: . ***.* .*.* **

H1/Cal09        ILGNPECESLSTASSWSYIVETPSSDNGTCYPGDFIDYEELREQLSSVSSFERFEIFPKT
H6              ILGNPKCDLLLGDQSWSYIVERPNAQNGICYPGVLNELEELKAFIGSGERVERFEMFPKS
                *****:*: *   .******* *.::** **** : : ***:  :.* . .****:***:

H1/Cal09        SSWPNHDSNKGVTAACP-HAGAKSFYKNLIWLVKKG-NSYPKLSKSYINDKGKEVLVLWG
H6              -TWAGVDTSRGVTNACPSYTIDSSFYRNLVWIVKTDSATYPVIKGTYNNTGTQPILYFWG
                 :*.. *:.:*** *** ::  .***:**:*:**..  :** :. :* *   : :* :**

H1/Cal09        IHHPSTSADQQSLYQNADTYVFVGSSRYSKKFKPEIAIRPKVRDQEGRMNYYWTLVEPGD
H6              VHHPLDTTVQDNLYGSGDKYVRMGTESMNFAKSPEIAARPAVNGQRSRIDYYWSVLRPGE
                :***  :: *:.** ..*.** :*:.  .   .**** ** *..*..*::***:::.**:

H1/Cal09        KITFEATGNLVVPRYAFAMERNAG-SGIIISDTPVHDCNTTCQTPKGAINTSLPFQNIHP
H6              TLNVESNGNLIAPWYAYKFVSTNKKGAVFKSDLPIENCDATCQTITGVLRTNKTFQNVSP
                .:..*:.***:.* **: :  .   ..:: ** *:.:*::**** .*.:.*. .***: *
                                              ⬇
H1/Cal09        ITIGKCPKYVKSTKLRLATGLRNIPSIQSRGLFGAIAGFIEGGWTGMVDGWYGYHHQNEQ
H6              LWIGECPKYVKSESLRLATGLRNVPQIATRGIFGAIAGFIEGGWTGMIDGWYGYHHENSQ
                : **:******* .*********:*.* :**:***************:********:*.*

H1/Cal09        GSGYAADLKSTQNAIDEITNKVNSVIEKMNTQFTAVGKEFNHLEKRIENLNKKVDDGFLD
H6              GSGYAADRESTQKAIDGITNKVNSIINKMNTQFEAVDHEFSNLERRIGNLNKRMEDGFLD
                ******* :***:*** *******:*:****** **.:**.:**:** ****:::*****

H1/Cal09        IWTYNAELLVLLENERTLDYHDSNVKNLYEKVRSQLKNNAKEIGNGCFEFYHKCDNTCME
H6              VWTYNAELLVLLENERTLDLHDANVKNLYEKVKSQLRDNANDLGNGCFEFWHKCDNECME
                :****************** **:*********:***::**:::*******:***** ***

H1/Cal09        SVKNGTYDYPKYSEEA
H6              SVKNGTYDYPKYQKES
                ************.:*:
```

**Supplementary Fig. 7 | Sequence alignment of HA ectodomains from H6 and H1/Cal09.** The boundary of the HA1 and HA2 chains is marked with an arrow. The amino acid sites that are computationally optimized are highlighted with red boxes.

```
H1/Cal09          DTLCIGYHANNSTDTVDTVLEKNVTVTHSVNLLEDKHNGKLCKLRGVAPLHLGKCNIAGW
H8                DRICIGYQSNNSTDTVNTLIEQNVPVTQTMELVETEKHPAYCNTDLGAPLELRDCKIEAV
                  * :****::*******:*::*:**.**::::*:* :::   *:    ***.* .*:* .

H1/Cal09          ILGNPECESLSTASSWSYIVETPSSDNGTCYPGDFIDYEELREQLSSVSSFERFEIFPKT
H8                IYGNPKCDIHLKDQGWSYIVERPSAPEGMCYPGSVENLEELRFVFSSAASYKRIRLFDYS
                  * ***:*:   . ..****** **: :* ****.. : ****  :**.:*::*:.:*  :

H1/Cal09          SSWPNHDSNKGVTAACPHAGAKSFYKNLIWLVKKG-NSYPKLSKSYINDKGKEVLVLWGI
H8                R--WNVTRSGTSKACNASTGGQSFYRSINWLTKKKPDTYDFNEGAYVNNEDGDIIFLWGI
                      *   .   .*. . :*.:***:.: **.**  ::*   . :*:*::. :::.****

H1/Cal09          HHPSTSADQQSLYQNADTYVFVGSSRYSKKFKPEIAIRPKVRDQEGRMNYYWTLVEPGDK
H8                HHPPDTKEQTTLYKNANTLSSVTTNTINRSFQPNIGPRPLVRGQQGRMDYYWGILKRGET
                  ***. : :* :**:**:*   * :.  .:.*:*:*. ** **.*:***:*** ::: *:.

H1/Cal09          ITFEATGNLVVPRYAFAMERNAGSGIIIS-DTPVHDCNTTCQTPKGAINTSLPFQNIHPI
H8                LKIRTNGNLIAPEFGYLLKGESYGRIIQNEDIPIGNCNTKCQTYAGAINSSKPFQNASRH
                  :.:.:.***:.*.:.: :: :: . ** . * *: :***.***  ****:* ****

H1/Cal09          TIGKCPKYVKSTKLRLATGLRNIPSIQSRGLFGAIAGFIEGGWTGMVDGWYGYHHQNEQG
H8                YMGECPKYVKKASLRLAVGLRNTPSVEPRGLFGAIAGFIEGGWSGMIDGWYGFHHSNSEG
                   :*:******.:.****.**** **::.***************:**:*****:**.*.:*

H1/Cal09          SGYAADLKSTQNAIDEITNKVNSVIEKMNTQFTAVGKEFNHLEKRIENLNKKVDDGFLDI
H8                TGMAADQKSTQEAIDKITNKVNNIVDKMNREFEVVNHEFSEVEKRINMINDKIDDQIEDL
                  :* *** ****:***:******.:::*** :* .*.:**..:****: :*.*:** : *:

H1/Cal09          WTYNAELLVLLENERTLDYHDSNVKNLYEKVRSQLKNNAKEIGNGCFEFYHKCDNTCMES
H8                WAYNAELLVLLENQKTLDEHDSNVKNLFDEVKRRLSANAIDAGNGCFDILHKCDNECMET
                  *:***********::*** ********:::*: :*. ** : *****:: ***** ***:

H1/Cal09          VKNGTYDYPKYSEEA
H8                IKNGTYDHKEYEEEA
                  :******: :*.***
```

**Supplementary Fig. 8 | Sequence alignment of HA ectodomains from H8 and H1/Cal09.** The boundary of the HA1 and HA2 chains is marked with an arrow. The amino acid sites that are computationally optimized are highlighted with red boxes.

```
H1/Cal09        DTLCIGYHANNSTDTVDTVLEKNVTVTHSVNLLEDKHNGKLCKLRGVAPLHLGKCNIAGW
H9              DKICIGYQTNNSTETVNTLSEQNVPVTQVEELVHGGIDPILCGTELGSPLVLDDCSLEGL
                *.:****::****:**:*: *:**.**:  :*:..  :  **  .  :** *..*.: *

H1/Cal09        ILGNPECESLSTASSWSYIVETPSSDNGTCYPGDFIDYEELREQLSSVSSFERFEIFPKT
H9              ILGNPKCDLYLNGREWSYIVERPKEMEGVCYPGSIENQEELRSLFSSIKKYERVKMFDFT
                *****:*:   .. .****** *.. :*.****.: : ****. :**:..:**.::*  *

H1/Cal09        SSWPNHDSNKGVTAACPHAGAKSFYKNLIWLVKKGNSYPKLSKSYINDKGKEVLVLWGIH
H9              KWNVTYTG--TSKACNNTSNQGSFYRSMRWLTLKSGQFPVQTDEYKNTRDSDIVFTWAIH
                .   .: .    .*.   :.  ***:.: **. *...:*  :..* * :..::.  *.**

H1/Cal09        HPSTSADQQSLYQNADTYVFVGSSRYSKKFKPEIAIRPKVRDQEGRMNYYWTLVEPGDKI
H9              HPPTSDEQVKLYKNPDTLSSVTTDEINRSFKPNIGPRPLVRGQQGRMDYYWAVLKPGQTV
                **.** :* .**:*.**   * :.. .:.***:*. ** **.*:***:***::::**:.:

H1/Cal09        TFEATGNLVVPRYAFAMERNAGSGIIISDTPVHDCNTTCQTPKGAINTSLPFQNIHPITI
H9              KIQTNGNLIAPEYGHLITGKSHGRILKNNLPMGQCVTECQLNEGVMNTSKPFQNTSKHYI
                .:::.***:.*.*.. :  :: . *: .: *: :* * **  :*.:*** ****     *

H1/Cal09        GKCPKYVKSTKLRLATGLRNIPSIQSRGLFGAIAGFIEGGWTGMVDGWYGYHHQNEQGSG
H9              GKCPKYIPSGSLKLAIGLRNVPQVQDRGLFGAIAGFIEGGWPGLVAGWYGFQHQNAEGTG
                ******: * .*:** ****:*.:*.***************.*:* ****::*** :*:*

H1/Cal09        YAADLKSTQNAIDEITNKVNSVIEKMNTQFTAVGKEFNHLEKRIENLNKKVDDGFLDIWT
H9              IAADRDSTQRAIDNMQNKLNNVIDKMNKQFEVVNHEFSEVESRINMINSKIDDQITDIWA
                 *** .***.***:: **:*.**:***.** .*.:**..:*.**: :*.*:** : ***:

H1/Cal09        YNAELLVLLENERTLDYHDSNVKNLYEKVRSQLKNNAKEIGNGCFEFYHKCDNTCMESVK
H9              YNAELLVLLENQKTLDEHDANVRNLHDRVRRVLRENAIDTGDGCFEILHKCDNNCMDTIR
                ***********::*** **:**:**:::**  *::** : *:****: *****.**::::

H1/Cal09        NGTYDYPKYSEEA
H9              NGTYNHKEYEEES
                ****:: :*.**:
```

**Supplementary Fig. 9 | Sequence alignment of HA ectodomains from H9 and H1/Cal09.** The boundary of the HA1 and HA2 chains is marked with an arrow. The amino acid sites that are computationally optimized are highlighted with red boxes.

```
H1/Cal09        DTLCIGYHANNSTDTVDTVLEKNVTVTHSVNLLEDKHNGKLCKLRGVAPLHLGKCNIAGW
H13             DRICVGYLSTNSSERVDTLLENGVPVTSSIDLIETNHTGTYCSLNGVSPVHLGDCSFEGW
                * :*:** :.**:: ***:**:.*.** *::*:* :*.*. *.*.**:*:***.*.: **

H1/Cal09        ILGNPECESLSTASSWSYIVETPSSDNGTCYPGDFIDYEELREQLSSVSSFERFEIFPKT
H13             IVGNPACTSNFGIREWSYLIEDPAAPHGLCYPGELNNNGELRHLFSGIRSFSRTELIPPT
                *:*** * *     .***::* *:: :* ****:: :  ***. :*.: **.* *::* *

H1/Cal09        SSWPNHDSNKGVTAACPHAGAKSFYKNLIWLVKKGNSYPKLSKSYINDKGKEVLVLWGIH
H13             SWGEVLDG--TTSACRDNTGTNSFYRNLVWFIKKNNRYPVISKTYNNTTGRDVLVLWGIH
                *     *.   .:*.  ::*::***:**:*::**.* ** :**:* * .*::********

H1/Cal09        HPSTSADQQSLYQNADTYVFVGSSRYSKKFKPEIAIRPKVRDQEGRMNYYWTLVEPGDKI
H13             HPVSVDETKTLYVNSDPYTLVSTKSWSEKYKLETGVRPGYNGQRSWMKIYWSLIHPGEMI
                ** :  : ::** *:*.*.:*.:. :*:*:* * .:**  ..*.. *: **:*:.**: *

H1/Cal09        TFEATGNLVVPRYAFAMERNAGSGIIISDTPVHDCNTTCQTPKGAINTSLPFQNIHPITI
H13             TFESNGGFLAPRYGYIIEEYGKGRIFQSRIRMSRCNTKCQTSVGGINTNRTFQNIDKNAL
                ***:.*.::.***.: :*. . . *: *   :  ***.***. *.***. .****.  ::

H1/Cal09        GKCPKYVKSTKLRLATGLRNIPSIQSRGLFGAIAGFIEGGWTGMVDGWYGYHHQNEQGSG
H13             GDCPKYIKSGQLKLATGLRNVPAISNRGLFGAIAGFIEGGWPGLINGWYGFQHQNEQGTG
                *.****:** :*:*******:*:*..***************.*:::****::******:*

H1/Cal09        YAADLKSTQNAIDEITNKVNSVIEKMNTQFTAVGKEFNHLEKRIENLNKKVDDGFLDIWT
H13             IAADKESTQKAIDQITTKINNIIDKMNGNYDSIRGEFNQVEKRINMLADRIDDAVTDIWS
                 *** :***:***:**.*:*.:*:*** :: ::  ***::****: * .::**.. ***:

H1/Cal09        YNAELLVLLENERTLDYHDSNVKNLYEKVRSQLKNNAKEIGNGCFEFYHKCDNTCMESVK
H13             YNAKLLVLLENDKTLDMHDANVKNLHEQVRRELKDNAIDEGNGCFELLHKCNDSCMETIR
                ***:*******::*** **:*****:*:** :**:** : ******: ***:::***:::

H1/Cal09        NGTYDYPKYSEEA
H13             NGTYDHTEYAEES
                *****:.:*:**:
```

**Supplementary Fig. 10 | Sequence alignment of HA ectodomains from H13 and H1/Cal09.** The boundary of the HA1 and HA2 chains is marked with an arrow. The amino acid sites that are computationally optimized are highlighted with red boxes.

```
H1/Cal09        DTLCIGYHANNSTDTVDTVLEKNVTVTHSVNLLEDKHNGKLCKLRGVAPLHLGKCNIAGW
H18             DQICIGYHSNNSTQTVNTLLESNVPVTSSHSILEKEHNGLLCKLKGKAPLDLIDCSLPAW
                * :*****:****:**:*:**.**.** * .:**.:*** ****:* ***.* .*.:..*

H1/Cal09        ILGNPECESLSTASSWSYIVETPSSDNGTCYPGDFIDYEELREQLSSVSSFERFEIFPKT
H18             LMGNPKCDELLTASEWAYIKEDPEPENGICFPGDFDSLEDLILLVSNTDHFRKEKIIDMT
                ::***:*:.* ***.*:** * *..:** *:**** . *:*   :*... *.: :*:  *

H1/Cal09        SSWPNHDSNKGVTAACPHAG-AKSFYKNLIWLVKKGNSYPKLSKSYINDKGKEVLVLWGI
H18             RFSDVTTNN--VDSACPYDTNGASFYRNLNWVQQ--NKGKQLIFHYQNSENNPLLIIWGV
                       .*  * :***:   . ***:** *: :  *.  :*   * *.:.: :*::**:

H1/Cal09        HHPSTSADQQSLYQNADTYVFVGSSRYSKKFKPEIAIRPKVRDQEGRMNYYWTLVEPGDK
H18             HQTSNAAEQNTYYGSQTGSTTITIGEETNTYPLVISESSILNGHSDRINYFWGVVNPNQN
                *:.*.:*:*:: * .    . :  .. ::.:   *:  . :..:..*:**:* :*:*.::

H1/Cal09        ITFEATGNLVVPRYAFAMERN-AGSGIIISDTPVHDCNTTCQTPKGAINTSLPFQNIHPI
H18             FSIVSTGNFIWPEYGYFFQKTTNISGIIKSSEKISDCDTICQTKIGAINSTLPFQNIHQN
                ::: :***:: *.*.: :::.   **** *.  : **:* ***  ****::*******

H1/Cal09        TIGKCPKYVKSTKLRLATGLRNIPSIQSRGLFGAIAGFIEGGWTGMVDGWYGYHHQNEQG
H18             AIGDCPKYVKAQELVLATGLRNNPIKETRGLFGAIAGFIEGGWQGLIDGWYGYHHQNSEG
                :**.******: :* ******* *  ::*************** *::**********.:*

H1/Cal09        SGYAADLKSTQNAIDEITNKVNSVIEKMNTQFTAVGKEFNHLEKRIENLNKKVDDGFLDI
H18             SGYAADKEATQKAVDAITTKVNNIIDKMNTQFESTAKEFNKIEMRIKHLSDRVDDGFLDV
                ****** ::**:*:* **.***.:*:****** :..****::* **::*..:*******:

H1/Cal09        WTYNAELLVLLENERTLDYHDSNVKNLYEKVRSQLKNNAKEIGNGCFEFYHKCDNTCMES
H18             WSYNAELLVLLENERTLDFHDANVNNLYQKVKVQLKDNAIDMGNGCFKILHKCNNTCMDD
                *:****************:**:**:***:**: ***:** ::*****:: ***:****:.

H1/Cal09        VKNGTYDYPKYSEEA
H18             IKNGTYNYYEYRKES
                :*****:* :* :*:
```

**Supplementary Fig. 11 | Sequence alignment of HA ectodomains from H18 and H1/Cal09.** The boundary of the HA1 and HA2 chains is marked with an arrow. The amino acid sites that are computationally optimized are highlighted with red boxes.

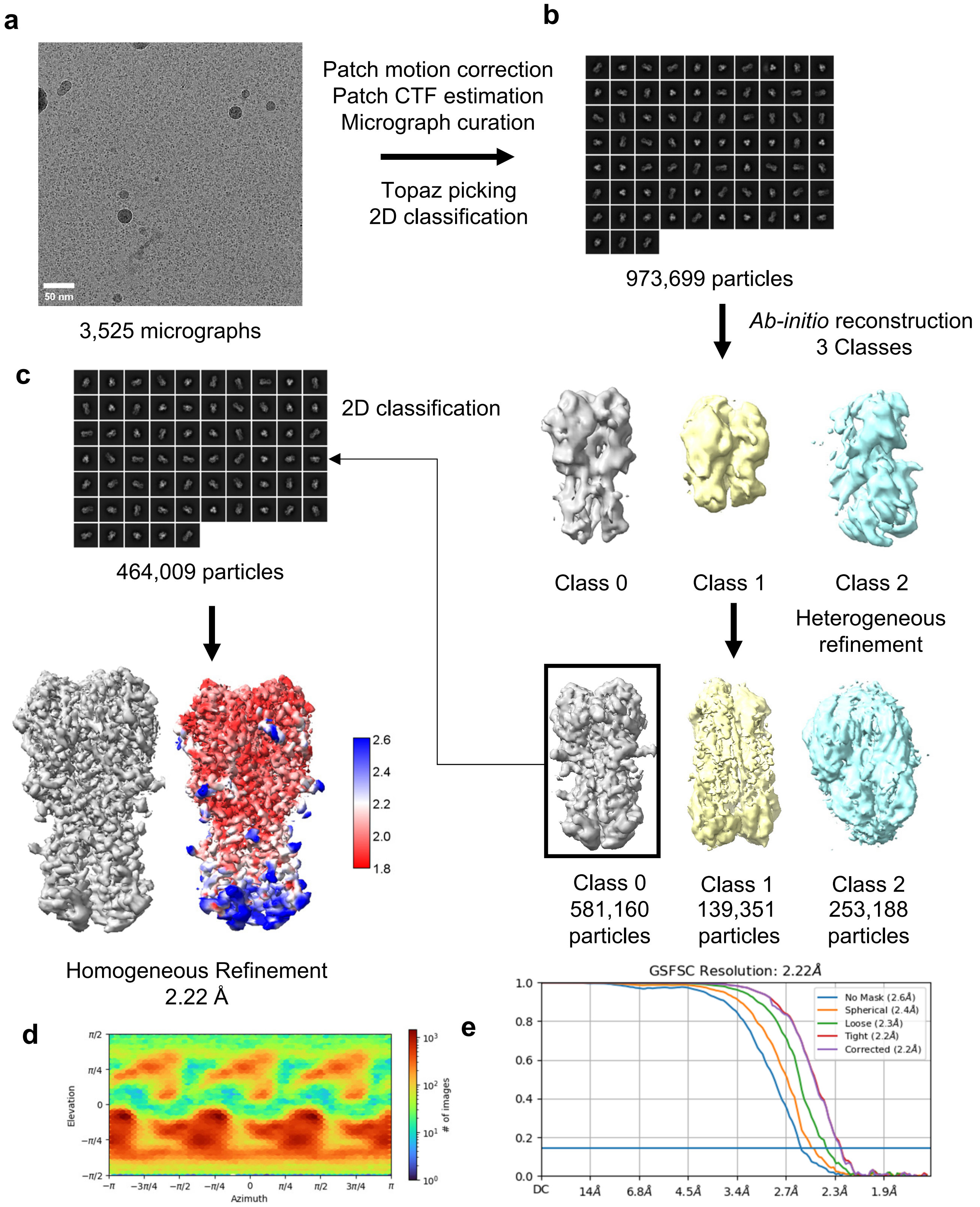

**Supplementary Fig. 12 | Cryo-EM data processing of the H1/Cal09-mut. a** A representative micrograph image. **b** Selected 2D class averages. **c** Summary of cryo-EM data processing. **d** Orientation distribution of the particles used for 3D electron density reconstruction. **e** Map FSC curve

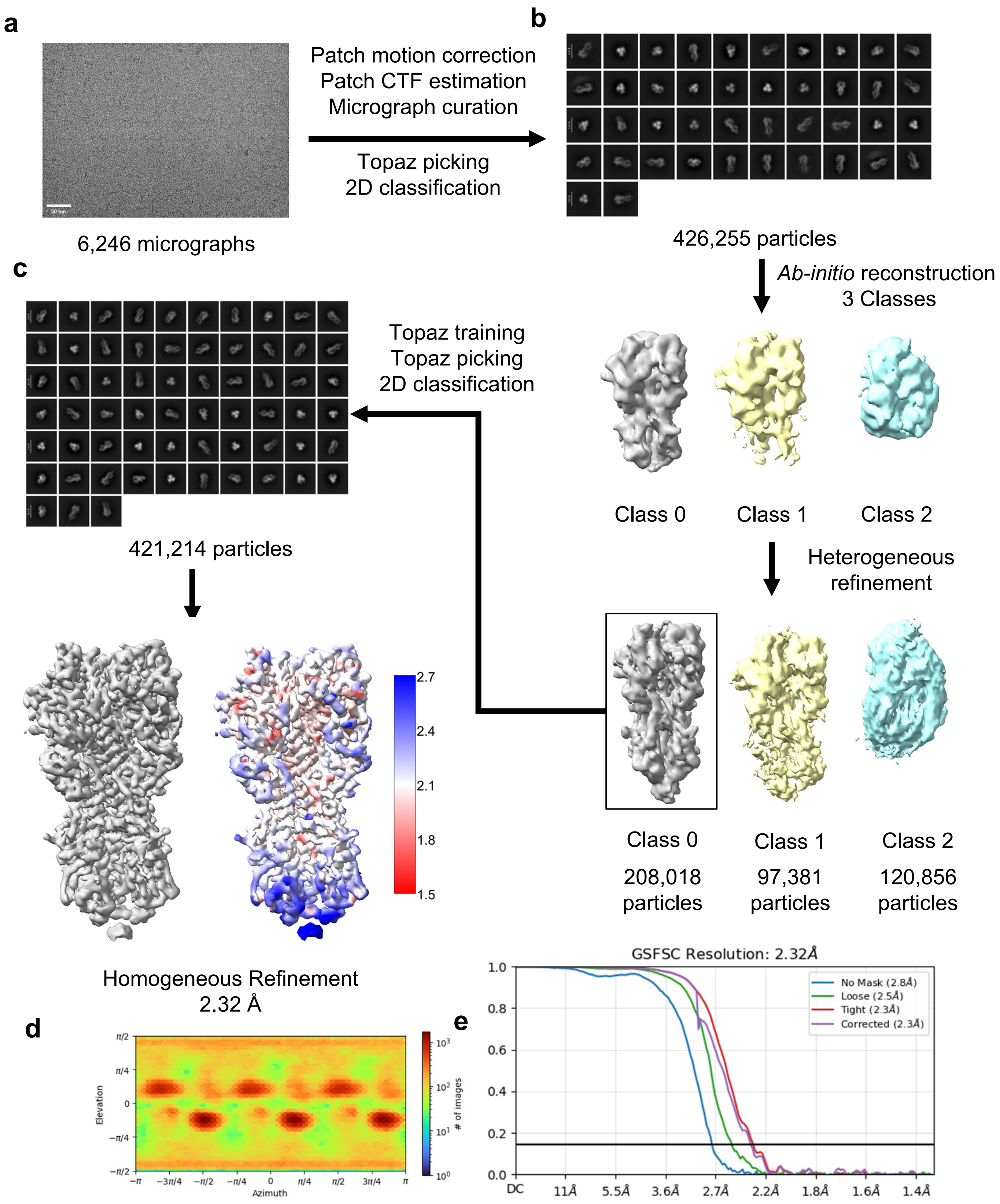

**Supplementary Fig. 13 | Cryo-EM data processing of the H1/Mal07-mut. a** A representative micrograph image. **b** Selected 2D class averages. **c** Summary of cryo-EM data processing. **d** Orientation distribution of the particles used for 3D electron density reconstruction. **e** Map FSC curve

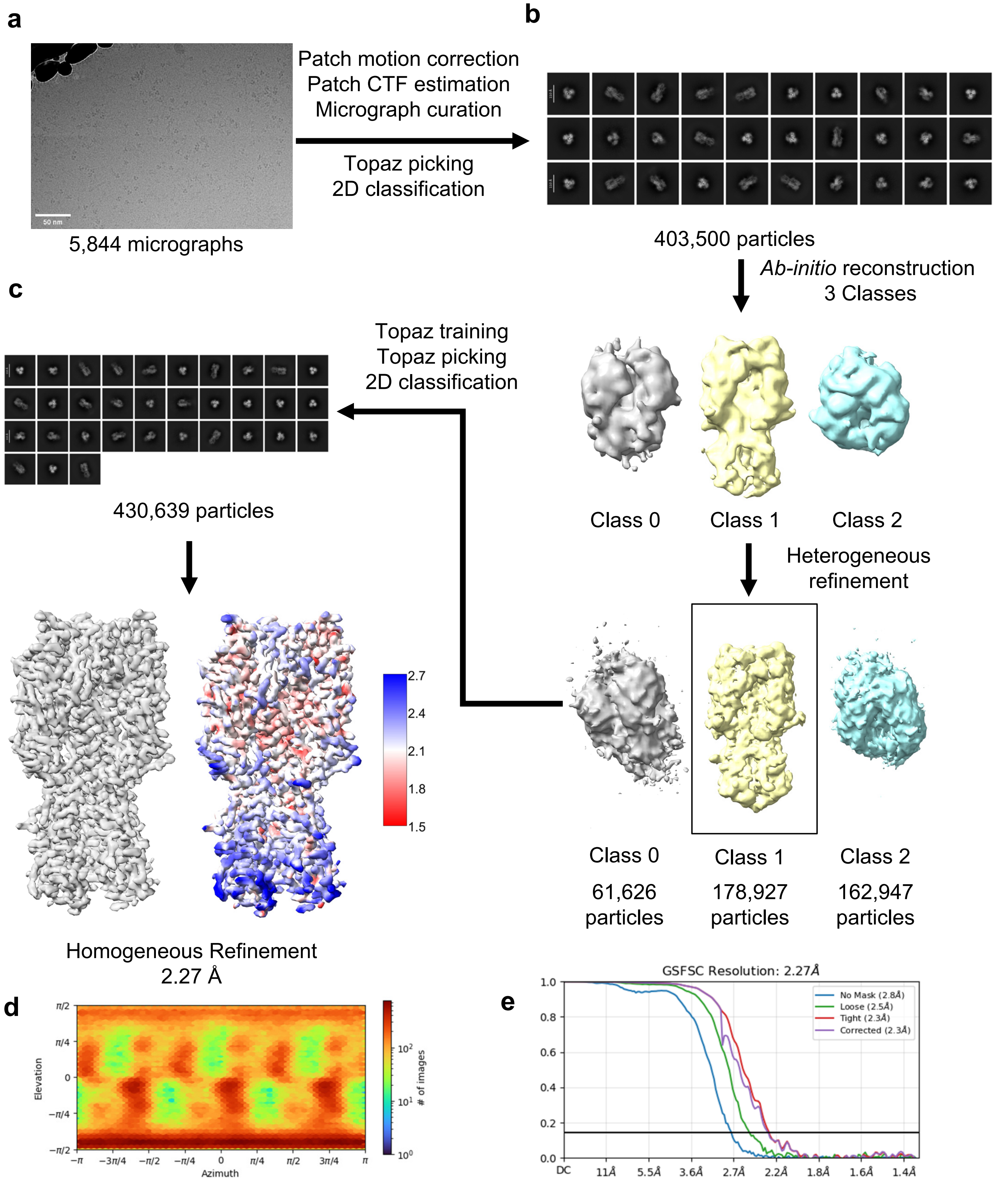

**Supplementary Fig. 14 | Cryo-EM data processing of the H9/HK98-mut. a** A representative micrograph image. **b** Selected 2D class averages. **c** Summary of cryo-EM data processing. **d** Orientation distribution of the particles used for 3D electron density reconstruction. **e** Map FSC curve

**Supplementary Table 1 | The purified H1/California/2009 HA protein sequences used for the structural and stability analyses**

| Protein name | Protein sequences |
|---|---|
| H1/Cal09-WT | DTLCIGYHANNSTDTVDTVLEKNVTVTHSVNLLEDKHNGKLCKLRGVAPLHLGKCNIA<br>GWILGNPECESLSTASSWSYIVETPSSDNGTCYPGDFIDYEELREQLSSVSSFERFEI<br>FPKTSSWPNHDSNKGVTAACPHAGAKSFYKNLIWLVKKGNSYPKLSKSYINDKGKEV<br>LVLWGIHHPSTSADQQSLYQNADTYVFVGSSRYSKKFKPEIAIRPKVRDQEGRMNYY<br>WTLVEPGDKITFEATGNLVVPRYAFAMERNAGSGIIISDTPVHDCNTTCQTPKGAINTS<br>LPFQNIHPITIGKCPKYVKSTKLRLATGLRNIPSIQSRGLFGAIAGFIEGGWTGMVDGW<br>YGYHHQNEQGSGYAADLKSTQNAIDEITNKVNSVIEKMNTQFTAVGKEFNHLEKRIEN<br>LNKKVDDGFLDIWTYNAELLVLLENERTLDYHDSNVKNLYEKVRSQLKNNAKEIGNGC<br>FEFYHKCDNTCMESVKNGTYDYPKYSEEAKLNREEIDSNSLEVLFQ |
| H1/Cal09-WT-tri | DTLCIGYHANNSTDTVDTVLEKNVTVTHSVNLLEDKHNGKLCKLRGVAPLHLGKCNIA<br>GWILGNPECESLSTASSWSYIVETPSSDNGTCYPGDFIDYEELREQLSSVSSFERFEI<br>FPKTSSWPNHDSNKGVTAACPHAGAKSFYKNLIWLVKKGNSYPKLSKSYINDKGKEV<br>LVLWGIHHPSTSADQQSLYQNADTYVFVGSSRYSKKFKPEIAIRPKVRDQEGRMNYY<br>WTLVEPGDKITFEATGNLVVPRYAFAMERNAGSGIIISDTPVHDCNTTCQTPKGAINTS<br>LPFQNIHPITIGKCPKYVKSTKLRLATGLRNIPSIQSRGLFGAIAGFIEGGWTGMVDGW<br>YGYHHQNEQGSGYAADLKSTQNAIDEITNKVNSVIEKMNTQFTAVGKEFNHLEKRIEN<br>LNKKVDDGFLDIWTYNAELLVLLENERTLDYHDSNVKNLYEKVRSQLKNNAKEIGNGC<br>FEFYHKCDNTCMESVKNGTYDYPKYSEEAKLNREEIDGSGYIPEAPRDGQAYVRKD<br>GEWVLLSTFLGNSLEVLFQ |
| H1/Cal09-mut | DTLCIGYHANNSTDTVDTVLEKNVTVTHSVNLLEDKHNGKLCKLRGVAPLHLGKCNIA<br>GWILGNPECESLSTASSWSYIVETPSSDNGTCYPGDFIDYEELREQLSSVSSFERFEI<br>FPKTSSWPNHDSNKGVTAACPHAGAKSFYKNLIWLVKKGNSYPKLSKSYINDKGKEV<br>LVLWGIHHPSTSADQQSLYQNADTYVFVGSSRYSKKFKPEIAIRPKVRDQEGRMNYY<br>WTLVEPGDKITFEATGNLVVPRYAFAMERNAGSGIIISDTPVHDCNTTCQTPKGAINTS<br>LPFQNIHPITIGKCPKYVKSTKLRLATGLRNIPSIQSRGLFGAIAGFIEGGWTGMVDGW<br>YGYHHQNEQGSGYAADLKSTQNAIDGITNKVNSVIEKMNTQFTAVGKEFNHLEKRIEN<br>LNKKVDDGFLDIWTYLAELLVLFLNERTLEYHDSNVKNLYEKVRSQLKNNAKEIGNGC<br>FEFYHKCDNTCMESVKNGTYDYPKYSEEAKLNREEIDSNSLEVLFQ |

Footnote: Non-native sequences including linkers, the Foldon sequence and PreScission protease sites are written in purple.

**Supplementary Table 2 | The purified H9/Hong Kong/98 HA protein sequences used for the structural and stability analyses**

| Protein name | Protein sequences |
|---|---|
| H9/HK98-WT | DKICIGYQSTNSTETVDTLTETNVPVTHAKELLHTSHNGMLCATNLGHPLILDTCTIEG<br>LIYGNPSCDLLLGGREWSYIVERPSAVNGMCYPGNVENLEELRSLFSSASSYQRIQIF<br>PDTIWNVSYSGTSSACSDSFYRSMRWLTQKNNAYPIQDAQYTNNRGKSILFMWGIN<br>HPPTDTVQTNLYTRTDTTTSVTTEDINRTFKPVIGPRPLVNGLHGRIDYYWSVLKPGQ<br>TLRVRSNGNLIAPWYGHILSGESHGRILKTDLNSGNCVVQCQTERGGLNTTLPFHNV<br>SKYAFGNCPKYVGVKSLKLAVGLRNVPARSSGLFGAIAGFIEGGWPGLVAGWYGFQH<br>SNDQGVGMAADSDSTQKAIDKITSKVNNIVDKMNKQYGIIDHEFSEIETRLNMINNKID<br>DQIQDIWTYNAELLVLLENQKTLDEHDANVNNLYNKVKRALGSNAMEDGKGCFELYH<br>KCDDQCMETIRNGTYNRRKYKEES SSGRLVPRGSHHHHHH |
| H9/HK98-WT-tri | DKICIGYQSTNSTETVDTLTETNVPVTHAKELLHTSHNGMLCATNLGHPLILDTCTIEG<br>LIYGNPSCDLLLGGREWSYIVERPSAVNGMCYPGNVENLEELRSLFSSASSYQRIQIF<br>PDTIWNVSYSGTSSACSDSFYRSMRWLTQKNNAYPIQDAQYTNNRGKSILFMWGIN<br>HPPTDTVQTNLYTRTDTTTSVTTEDINRTFKPVIGPRPLVNGLHGRIDYYWSVLKPGQ<br>TLRVRSNGNLIAPWYGHILSGESHGRILKTDLNSGNCVVQCQTERGGLNTTLPFHNV<br>SKYAFGNCPKYVGVKSLKLAVGLRNVPARSSGLFGAIAGFIEGGWPGLVAGWYGFQH<br>SNDQGVGMAADSDSTQKAIDKITSKVNNIVDKMNKQYGIIDHEFSEIETRLNMINNKID<br>DQIQDIWTYNAELLVLLENQKTLDEHDANVNNLYNKVKRALGSNAMEDGKGCFELYH<br>KCDDQCMETIRNGTYNRRKYKEESGSGYIPEAPRDGQAYVRKDGEWVLLSTFLGRS<br>SGRLVPRGSHHHHHH |
| H9/HK98-mut | DKICIGYQSTNSTETVDTLTETNVPVTHAKELLHTSHNGMLCATNLGHPLILDTCTIEG<br>LIYGNPSCDLLLGGREWSYIVERPSAVNGMCYPGNVENLEELRSLFSSASSYQRIQIF<br>PDTIWNVSYSGTSSACSDSFYRSMRWLTQKNNAYPIQDAQYTNNRGKSILFMWGIN<br>HPPTDTVQTNLYTRTDTTTSVTTEDINRTFKPVIGPRPLVNGLHGRIDYYWSVLKPGQ<br>TLRVRSNGNLIAPWYGHILSGESHGRILKTDLNSGNCVVQCQTERGGLNTTLPFHNV<br>SKYAFGNCPKYVGVKSLKLAVGLRNVPARSSGLFGAIAGFIEGGWPGLVAGWYGFQ<br>HSNDQGVGMAADSDSTQKAIDKITSKVNNIVDKMNKQYGIIDHEFSEIETRLNMINNKI<br>DDQIQDIWTYLAELLVLLLNQKTLDEHDANVNNLYNKVKRALGSNAMEDGKGCFELY<br>HKCDDQCMETIRNGTYNRRKYKEES SSGRLVPRGSHHHHHH |
| H9/HK98-mut-tri | DKICIGYQSTNSTETVDTLTETNVPVTHAKELLHTSHNGMLCATNLGHPLILDTCTIEG<br>LIYGNPSCDLLLGGREWSYIVERPSAVNGMCYPGNVENLEELRSLFSSASSYQRIQIF<br>PDTIWNVSYSGTSSACSDSFYRSMRWLTQKNNAYPIQDAQYTNNRGKSILFMWGIN<br>HPPTDTVQTNLYTRTDTTTSVTTEDINRTFKPVIGPRPLVNGLHGRIDYYWSVLKPGQ<br>TLRVRSNGNLIAPWYGHILSGESHGRILKTDLNSGNCVVQCQTERGGLNTTLPFHNV<br>SKYAFGNCPKYVGVKSLKLAVGLRNVPARSSGLFGAIAGFIEGGWPGLVAGWYGFQH<br>SNDQGVGMAADSDSTQKAIDKITSKVNNIVDKMNKQYGIIDHEFSEIETRLNMINNKID<br>DQIQDIWTYLAELLVLLLNQKTLDEHDANVNNLYNKVKRALGSNAMEDGKGCFELYH<br>KCDDQCMETIRNGTYNRRKYKEESGSGYIPEAPRDGQAYVRKDGEWVLLSTFLGRS<br>SGRLVPRGSHHHHHH |

Footnote: Non-native sequences including linkers, the Foldon sequences, histidine tags and thrombin sites are written in purple.

**Supplementary Table 3 | The purified H1/Malaysia/2007 HA protein sequences used for the structural and stability analyses**

| Protein name | Protein sequences |
|---|---|
| H1/Mal07-WT | DTICIGYHANNSTDTVDTVLEKNVTVTHSVNLLEDSHNGKLCLLKGIAPLQLGNCSVA<br>GWILGNPECELLISRESWSYIVEKPNPENGTCYPGHFADYEELREQLSSVSSFERFEI<br>FPKESSWPNHTTTGVSASCSHNGESSFYKNLLWLTGKNGLYPNLSKSYANNKEKEVL<br>VLWGVHHPPNIGDQRALYHTENAYVSVVSSHYSRKFTPEIAKRPKVRDQEGRINYYW<br>TLLEPGDTIIFEANGNLIAPRYAFALSRGFGSGIINSNAPMDECDAKCQTPQGAINSSL<br>PFQNVHPVTIGECPKYVRSAKLRMVTGLRNIPSIQSRGLFGAIAGFIEGGWTGMVDG<br>WYGYHHQNEQGSGYAADQKSTQNAINGITNKVNSVIEKMNTQFTAVGKEFNKLERR<br>MENLNKKVDDGFIDIWTYNAELLVLLENERTLDFHDSNVKNLYEKVKSQLKNNAKEIG<br>NGCFEFYHKCNDECMESVKNGTYDYPKYSEESKLNREKIDSSGRLVPRGSHHHHHH |
| H1/Mal07-WT-tri | DTICIGYHANNSTDTVDTVLEKNVTVTHSVNLLEDSHNGKLCLLKGIAPLQLGNCSVA<br>GWILGNPECELLISRESWSYIVEKPNPENGTCYPGHFADYEELREQLSSVSSFERFEI<br>FPKESSWPNHTTTGVSASCSHNGESSFYKNLLWLTGKNGLYPNLSKSYANNKEKEVL<br>VLWGVHHPPNIGDQRALYHTENAYVSVVSSHYSRKFTPEIAKRPKVRDQEGRINYYW<br>TLLEPGDTIIFEANGNLIAPRYAFALSRGFGSGIINSNAPMDECDAKCQTPQGAINSSL<br>PFQNVHPVTIGECPKYVRSAKLRMVTGLRNIPSIQSRGLFGAIAGFIEGGWTGMVDG<br>WYGYHHQNEQGSGYAADQKSTQNAINGITNKVNSVIEKMNTQFTAVGKEFNKLERR<br>MENLNKKVDDGFIDIWTYNAELLVLLENERTLDFHDSNVKNLYEKVKSQLKNNAKEIG<br>NGCFEFYHKCNDECMESVKNGTYDYPKYSEESKLNREKIDGSGYIPEAPRDGQAYV<br>RKDGEWVLLSTFLGRSSGRLVPRGSHHHHHH |
| H1/Mal07-mut | DTICIGYHANNSTDTVDTVLEKNVTVTHSVNLLEDSHNGKLCLLKGIAPLQLGNCSVA<br>GWILGNPECELLISRESWSYIVEKPNPENGTCYPGHFADYEELREQLSSVSSFERFEI<br>FPKESSWPNHTTTGVSASCSHNGESSFYKNLLWLTGKNGLYPNLSKSYANNKEKEV<br>LVLWGVHHPPNIGDQRALYHTENAYVSVVSSHYSRKFTPEIAKRPKVRDQEGRINYY<br>WTLLEPGDTIIFEANGNLIAPRYAFALSRGFGSGIINSNAPMDECDAKCQTPQGAINSS<br>LPFQNVHPVTIGECPKYVRSAKLRMVTGLRNIPSIQSRGLFGAIAGFIEGGWTGMVDG<br>WYGYHHQNEQGSGYAADQKSTQNAINGITNKVNSVIEKMNTQFTAVGKEFNKLERR<br>MENLNKKVDDGFIDIWTYLAELLVLFLNERTLEFHDSNVKNLYEKVKSQLKNNAKEIG<br>NGCFEFYHKCNDECMESVKNGTYDYPKYSEESKLNREKIDSSGRLVPRGSHHHHH<br>H |
| H1/Mal07-mut-tri | DTICIGYHANNSTDTVDTVLEKNVTVTHSVNLLEDSHNGKLCLLKGIAPLQLGNCSVA<br>GWILGNPECELLISRESWSYIVEKPNPENGTCYPGHFADYEELREQLSSVSSFERFEI<br>FPKESSWPNHTTTGVSASCSHNGESSFYKNLLWLTGKNGLYPNLSKSYANNKEKEV<br>LVLWGVHHPPNIGDQRALYHTENAYVSVVSSHYSRKFTPEIAKRPKVRDQEGRINYY<br>WTLLEPGDTIIFEANGNLIAPRYAFALSRGFGSGIINSNAPMDECDAKCQTPQGAINSS<br>LPFQNVHPVTIGECPKYVRSAKLRMVTGLRNIPSIQSRGLFGAIAGFIEGGWTGMVDG<br>WYGYHHQNEQGSGYAADQKSTQNAINGITNKVNSVIEKMNTQFTAVGKEFNKLERR<br>MENLNKKVDDGFIDIWTYLAELLVLFLNERTLEFHDSNVKNLYEKVKSQLKNNAKEIG<br>NGCFEFYHKCNDECMESVKNGTYDYPKYSEESKLNREKIDGSGYIPEAPRDGQAYV<br>RKDGEWVLLSTFLGRSSGRLVPRGSHHHHHH |

Footnote: Non-native sequences including linkers, the Foldon sequences, histidine tags and thrombin sites are written in purple.

**Supplementary Table 4 | The purified H1/Malaysia/2007 HA proteins with three mutations used for the structural and stability analyses**

| Protein name | Protein sequences |
|---|---|
| H1/Mal07-(rev-95L) | DTICIGYHANNSTDTVDTVLEKNVTVTHSVNLLEDSHNGKLCLLKGIAPLQLGNCSVA<br>GWILGNPECELLISRESWSYIVEKPNPENGTCYPGHFADYEELREQLSSVSSFERFEI<br>FPKESSWPNHTTTGVSASCSHNGESSFYKNLLWLTGKNGLYPNLSKSYANNKEKEVL<br>VLWGVHHPPNIGDQRALYHTENAYVSVVSSHYSRKFTPEIAKRPKVRDQEGRINYYW<br>TLLEPGDTIIFEANGNLIAPRYAFALSRGFGSGIINSNAPMDECDAKCQTPQGAINSSL<br>PFQNVHPVTIGECPKYVRSAKLRMVTGLRNIPSIQSRGLFGAIAGFIEGGWTGMVDG<br>WYGYHHQNEQGSGYAADQKSTQNAINGITNKVNSVIEKMNTQFTAVGKEFNKLERR<br>MENLNKKVDDGFIDIWTYNAELLVLFLNERTLEFHDSNVKNLYEKVKSQLKNNAKEIG<br>NGCFEFYHKCNDECMESVKNGTYDYPKYSEESKLNREKIDSSGRLVPRGSHHHHHH |
| H1/Mal07-(rev-102F) | DTICIGYHANNSTDTVDTVLEKNVTVTHSVNLLEDSHNGKLCLLKGIAPLQLGNCSVA<br>GWILGNPECELLISRESWSYIVEKPNPENGTCYPGHFADYEELREQLSSVSSFERFEI<br>FPKESSWPNHTTTGVSASCSHNGESSFYKNLLWLTGKNGLYPNLSKSYANNKEKEVL<br>VLWGVHHPPNIGDQRALYHTENAYVSVVSSHYSRKFTPEIAKRPKVRDQEGRINYYW<br>TLLEPGDTIIFEANGNLIAPRYAFALSRGFGSGIINSNAPMDECDAKCQTPQGAINSSL<br>PFQNVHPVTIGECPKYVRSAKLRMVTGLRNIPSIQSRGLFGAIAGFIEGGWTGMVDG<br>WYGYHHQNEQGSGYAADQKSTQNAINGITNKVNSVIEKMNTQFTAVGKEFNKLERR<br>MENLNKKVDDGFIDIWTYLAELLVLLLNERTLEFHDSNVKNLYEKVKSQLKNNAKEIGN<br>GCFEFYHKCNDECMESVKNGTYDYPKYSEESKLNREKIDSSGRLVPRGSHHHHHH |
| H1/Mal07-(rev-103L) | DTICIGYHANNSTDTVDTVLEKNVTVTHSVNLLEDSHNGKLCLLKGIAPLQLGNCSVA<br>GWILGNPECELLISRESWSYIVEKPNPENGTCYPGHFADYEELREQLSSVSSFERFEI<br>FPKESSWPNHTTTGVSASCSHNGESSFYKNLLWLTGKNGLYPNLSKSYANNKEKEVL<br>VLWGVHHPPNIGDQRALYHTENAYVSVVSSHYSRKFTPEIAKRPKVRDQEGRINYYW<br>TLLEPGDTIIFEANGNLIAPRYAFALSRGFGSGIINSNAPMDECDAKCQTPQGAINSSL<br>PFQNVHPVTIGECPKYVRSAKLRMVTGLRNIPSIQSRGLFGAIAGFIEGGWTGMVDG<br>WYGYHHQNEQGSGYAADQKSTQNAINGITNKVNSVIEKMNTQFTAVGKEFNKLERR<br>MENLNKKVDDGFIDIWTYLAELLVLFENERTLEFHDSNVKNLYEKVKSQLKNNAKEIG<br>NGCFEFYHKCNDECMESVKNGTYDYPKYSEESKLNREKIDSSGRLVPRGSHHHHHH |
| H1/Mal07-(rev-109E) | DTICIGYHANNSTDTVDTVLEKNVTVTHSVNLLEDSHNGKLCLLKGIAPLQLGNCSVA<br>GWILGNPECELLISRESWSYIVEKPNPENGTCYPGHFADYEELREQLSSVSSFERFEI<br>FPKESSWPNHTTTGVSASCSHNGESSFYKNLLWLTGKNGLYPNLSKSYANNKEKEVL<br>VLWGVHHPPNIGDQRALYHTENAYVSVVSSHYSRKFTPEIAKRPKVRDQEGRINYYW<br>TLLEPGDTIIFEANGNLIAPRYAFALSRGFGSGIINSNAPMDECDAKCQTPQGAINSSL<br>PFQNVHPVTIGECPKYVRSAKLRMVTGLRNIPSIQSRGLFGAIAGFIEGGWTGMVDG<br>WYGYHHQNEQGSGYAADQKSTQNAINGITNKVNSVIEKMNTQFTAVGKEFNKLERR<br>MENLNKKVDDGFIDIWTYLAELLVLFLNERTLDFHDSNVKNLYEKVKSQLKNNAKEIG<br>NGCFEFYHKCNDECMESVKNGTYDYPKYSEESKLNREKIDSSGRLVPRGSHHHHHH |

Footnote: Non-native sequences including linkers, the Foldon sequences, histidine tags and thrombin sites are written in purple.

**Supplementary Table 5 | The purified H5 and H6 HA protein sequences used for the gel permeation chromatographic analyses**

| Protein name | Protein sequences |
|---|---|
| H5-WT | ADPDQICIGYHANNSTEQVDTIMEKNVTVTHAQDILEKTHNGKLCDLDGVKPLILRDC<br>SVAGWLLGNPMCDEFINVPEWSYIVEKANPVNDLCYPGDFNDYEELKHLLSRINHFE<br>KIQIIPKSSWSSHEASLGVSSACPYQGKSSFFRNVVWLIKKNSTYPTIKRSYNNTNQE<br>DLLVLWGIHHPNDAAEQTKLYQNPTTYISVGTSTLNQRLVPRIATRSKVNGQSGRMEF<br>FWTILKPNDAINFESNGNFIAPEYAYKIVKKGDSTIMKSELEYGNCNTKCQTPMGAINS<br>SMPFHNIHPLTIGECPKYVKSNRLVLATGLRNSPQRETRGLFGAIAGFIEGGWQGMV<br>DGWYGYHHSNEQGSGYAADKESTQKAIDGVTNKVNSIIDKMNTQFEAVGREFNNLE<br>RRIENLNKKMEDGFLDVWTYNAELLVLMENERTLDFHDSNVKNLYDKVRLQLRDNAK<br>ELGNGCFEFYHKCDNECMESVRNGTYDYPQYSEEAGGRLVPRGSHHHHHH |
| H5-mut | ADPDQICIGYHANNSTEQVDTIMEKNVTVTHAQDILEKTHNGKLCDLDGVKPLILRDC<br>SVAGWLLGNPMCDEFINVPEWSYIVEKANPVNDLCYPGDFNDYEELKHLLSRINHFE<br>KIQIIPKSSWSSHEASLGVSSACPYQGKSSFFRNVVWLIKKNSTYPTIKRSYNNTNQE<br>DLLVLWGIHHPNDAAEQTKLYQNPTTYISVGTSTLNQRLVPRIATRSKVNGQSGRMEF<br>FWTILKPNDAINFESNGNFIAPEYAYKIVKKGDSTIMKSELEYGNCNTKCQTPMGAINS<br>SMPFHNIHPLTIGECPKYVKSNRLVLATGLRNSPQRETRGLFGAIAGFIEGGWQGMV<br>DGWYGYHHSNEQGSGYAADKESTQKAIDYVTNKVNSIIDKMNTQFEAVGREFNNLE<br>RRIENLNKKMEDGFLDVWTYLAELLVLLLNERTLNFHDSNVKNLYDKVRLQLRDNAKE<br>LGNGCFEFYHKCDNECMESVRNGTYDYPQYSEEAGGRLVPRGSHHHHHH |
| H6-WT | ADPDKICIGYHANNSTTQVDTLLEKNVTVTHSVELLENQKEKRFCKIMNKAPLDLKDC<br>TIEGWILGNPKCDLLLGDQSWSYIVERPNAQNGICYPGVLNELEELKAFIGSGERVER<br>FEMFPKSTWAGVDTSRGVTNACPSYTIDSSFYRNLVWIVKTDSATYPVIKGTYNNTGT<br>QPILYFWGVHHPLDTTVQDNLYGSGDKYVRMGTESMNFAKSPEIAARPAVNGQRSRI<br>DYYWSVLRPGETLNVESNGNLIAPWYAYKFVSTNKKGAVFKSDLPIENCDATCQTITG<br>VLRTNKTFQNVSPLWIGECPKYVKSESLRLATGLRNVPQIATRGIFGAIAGFIEGGWTG<br>MIDGWYGYHHENSQGSGYAADRESTQKAIDGITNKVNSIINKMNTQFEAVDHEFSNL<br>ERRIGNLNKRMEDGFLDVWTYNAELLVLLENERTLDLHDANVKNLYEKVKSQLRDNA<br>NDLGNGCFEFWHKCDNECMESVKNGTYDYPKYQKESGGRLVPRGSHHHHHH |
| H6-mut | ADPDKICIGYHANNSTTQVDTLLEKNVTVTHSVELLENQKEKRFCKIMNKAPLDLKDC<br>TIEGWILGNPKCDLLLGDQSWSYIVERPNAQNGICYPGVLNELEELKAFIGSGERVER<br>FEMFPKSTWAGVDTSRGVTNACPSYTIDSSFYRNLVWIVKTDSATYPVIKGTYNNTGT<br>QPILYFWGVHHPLDTTVQDNLYGSGDKYVRMGTESMNFAKSPEIAARPAVNGQRSRI<br>DYYWSVLRPGETLNVESNGNLIAPWYAYKFVSTNKKGAVFKSDLPIENCDATCQTITG<br>VLRTNKTFQNVSPLWIGECPKYVKSESLRLATGLRNVPQIATRGIFGAIAGFIEGGWTG<br>MIDGWYGYHHENSQGSGYAADRESTQKAIDIITNKVNSIINKMNTQFEAVDHEFSNLE<br>RRIGNLNKRMEDGFLDVWTYLAELLVLLLNERTLALHDANVKNLYEKVKSQLRDNAN<br>DLGNGCFEFWHKCDNECMESVKNGTYDYPKYQKESGGRLVPRGSHHHHHH |

Footnote: Non-native sequences including linkers, the Foldon sequences, histidine tags and thrombin sites are written in purple.

**Supplementary Table 6 | The purified H8 and H12 HA protein sequences used for the gel permeation chromatographic analyses**

| Protein name | Protein sequences |
| --- | --- |
| H8-WT | ADPDRICIGYQSNNSTDTVNTLIEQNVPVTQTMELVETEKHPAYCNTDLGAPLELRDC<br>KIEAVIYGNPKCDIHLKDQGWSYIVERPSAPEGMCYPGSVENLEELRFVFSSAASYKR<br>IRLFDYSRWNVTRSGTSKACNASTGGQSFYRSINWLTKKKPDTYDFNEGAYVNNED<br>GDIIFLWGIHHPPDTKEQTTLYKNANTLSSVTTNTINRSFQPNIGPRPLVRGQQGRMD<br>YYWGILKRGETLKIRTNGNLIAPEFGYLLKGESYGRIIQNEDIPIGNCNTKCQTYAGAIN<br>SSKPFQNASRHYMGECPKYVKKASLRLAVGLRNTPSVEPRGLFGAIAGFIEGGWSG<br>MIDGWYGFHHSNSEGTGMAADQKSTQEAIDKITNKVNNIVDKMNREFEVVNHEFSE<br>VEKRINMINDKIDDQIEDLWAYNAELLVLLENQKTLDEHDSNVKNLFDEVKRRLSANAI<br>DAGNGCFDILHKCDNECMETIKNGTYDHKEYEEEAGGRLVPRGSHHHHHH |
| H8-mut | ADPDRICIGYQSNNSTDTVNTLIEQNVPVTQTMELVETEKHPAYCNTDLGAPLELRDC<br>KIEAVIYGNPKCDIHLKDQGWSYIVERPSAPEGMCYPGSVENLEELRFVFSSAASYKR<br>IRLFDYSRWNVTRSGTSKACNASTGGQSFYRSINWLTKKKPDTYDFNEGAYVNNED<br>GDIIFLWGIHHPPDTKEQTTLYKNANTLSSVTTNTINRSFQPNIGPRPLVRGQQGRMD<br>YYWGILKRGETLKIRTNGNLIAPEFGYLLKGESYGRIIQNEDIPIGNCNTKCQTYAGAIN<br>SSKPFQNASRHYMGECPKYVKKASLRLAVGLRNTPSVEPRGLFGAIAGFIEGGWSG<br>MIDGWYGFHHSNSEGTGMAADQKSTQEAIDKITNKVNNIVDKMNREFEVVNHEFSE<br>VEKRINMINDKIDDQIEDLWAYLAELLVLLLNQKTLSEHDSNVKNLFDEVKRRLSANAID<br>AGNGCFDILHKCDNECMETIKNGTYDHKEYEEEAGGRLVPRGSHHHHHH |
| H12-WT | ADPDKICIGYQTNNSTETVNTLSEQNVPVTQVEELVHGGIDPILCGTELGSPLVLDDCS<br>LEGLILGNPKCDLYLNGREWSYIVERPKEMEGVCYPGSIENQEELRSLFSSIKKYERV<br>KMFDFTKWNVTYTGTSKACNNTSNQGSFYRSMRWLTLKSGQFPVQTDEYKNTRDS<br>DIVFTWAIHHPPTSDEQVKLYKNPDTLSSVTTDEINRSFKPNIGPRPLVRGQQGRMDY<br>YWAVLKPGQTVKIQTNGNLIAPEYGHLITGKSHGRILKNNLPMGQCVTECQLNEGVM<br>NTSKPFQNTSKHYIGKCPKYIPSGSLKLAIGLRNVPQVQDRGLFGAIAGFIEGGWPGL<br>VAGWYGFQHQNAEGTGIAADRDSTQRAIDNMQNKLNNVIDKMNKQFEVVNHEFSEV<br>ESRINMINSKIDDQITDIWAYNAELLVLLENQKTLDEHDANVRNLHDRVRRVLRENAID<br>TGDGCFEILHKCDNNCMDTIRNGTYNHKEYEEESGGRLVPRGSHHHHHH |
| H12-mut | ADPDKICIGYQTNNSTETVNTLSEQNVPVTQVEELVHGGIDPILCGTELGSPLVLDDCS<br>LEGLILGNPKCDLYLNGREWSYIVERPKEMEGVCYPGSIENQEELRSLFSSIKKYERV<br>KMFDFTKWNVTYTGTSKACNNTSNQGSFYRSMRWLTLKSGQFPVQTDEYKNTRDS<br>DIVFTWAIHHPPTSDEQVKLYKNPDTLSSVTTDEINRSFKPNIGPRPLVRGQQGRMDY<br>YWAVLKPGQTVKIQTNGNLIAPEYGHLITGKSHGRILKNNLPMGQCVTECQLNEGVM<br>NTSKPFQNTSKHYIGKCPKYIPSGSLKLAIGLRNVPQVQDRGLFGAIAGFIEGGWPGL<br>VAGWYGFQHQNAEGTGIAADRDSTQRAIDAMQNKLNNVIDKMNKQFEVVNHEFSEV<br>ESRINMINSKIDDQITDIWAYLAELLVLLLNQKTLDEHDANVRNLHDRVRRVLRENAIDT<br>GDGCFEILHKCDNNCMDTIRNGTYNHKEYEEESGGRLVPRGSHHHHHH |

Footnote: Non-native sequences including linkers, histidine tags and thrombin sites are written in purple.

**Supplementary Table 7 | The purified H13 and H18 HA protein sequences used for the gel permeation chromatographic analyses**

| Protein name | Protein sequences |
|---|---|
| H13-WT | ADPDRICVGYLSTNSSERVDTLLENGVPVTSSIDLIETNHTGTYCSLNGVSPVHLGDC<br>SFEGWIVGNPACTSNFGIREWSYLIEDPAAPHGLCYPGELNNNGELRHLFSGIRSFSR<br>TELIPPTSWGEVLDGTTSACRDNTGTNSFYRNLVWFIKKNNRYPVISKTYNNTTGRD<br>VLVLWGIHHPVSVDETKTLYVNSDPYTLVSTKSWSEKYKLETGVRPGYNGQRSWMKI<br>YWSLIHPGEMITFESNGGFLAPRYGYIIEEYGKGRIFQSRIRMSRCNTKCQTSVGGIN<br>TNRTFQNIDKNALGDCPKYIKSGQLKLATGLRNVPAISNRGLFGAIAGFIEGGWPGLIN<br>GWYGFQHQNEQGTGIAADKESTQKAIDQITTKINNIIDKMNGNYDSIRGEFNQVEKRI<br>NMLADRIDDAVTDIWSYNAKLLVLLENDKTLDMHDANVKNLHEQVRRELKDNAIDEG<br>NGCFELLHKCNDSCMETIRNGTYDHTEYAEESGGRLVPRGSHHHHHH |
| H13-mut | ADPDRICVGYLSTNSSERVDTLLENGVPVTSSIDLIETNHTGTYCSLNGVSPVHLGDC<br>SFEGWIVGNPACTSNFGIREWSYLIEDPAAPHGLCYPGELNNNGELRHLFSGIRSFSR<br>TELIPPTSWGEVLDGTTSACRDNTGTNSFYRNLVWFIKKNNRYPVISKTYNNTTGRD<br>VLVLWGIHHPVSVDETKTLYVNSDPYTLVSTKSWSEKYKLETGVRPGYNGQRSWMKI<br>YWSLIHPGEMITFESNGGFLAPRYGYIIEEYGKGRIFQSRIRMSRCNTKCQTSVGGIN<br>TNRTFQNIDKNALGDCPKYIKSGQLKLATGLRNVPAISNRGLFGAIAGFIEGGWPGLIN<br>GWYGFQHQNEQGTGIAADKESTQKAIDYITTKINNIIDKMNGNYDSIRGEFNQVEKRI<br>NMLADRIDDAVTDIWSYLAKLLVLLLNDKTLAMHDANVKNLHEQVRRELKDNAIDEGN<br>GCFELLHKCNDSCMETIRNGTYDHTEYAEESGGRLVPRGSHHHHHH |
| H18-WT | ADPDQICIGYHSNNSTQTVNTLLESNVPVTSSHSILEKEHNGLLCKLKGKAPLDLIDCS<br>LPAWLMGNPKCDELLTASEWAYIKEDPEPENGICFPGDFDSLEDLILLVSNTDHFRKE<br>KIIDMTRFSDVTTNNVDSACPYDTNGASFYRNLNWVQQNKGKQLIFHYQNSENNPLL<br>IIWGVHQTSNAAEQNTYYGSQTGSTTITIGEETNTYPLVISESSILNGHSDRINYFWGV<br>VNPNQNFSIVSTGNFIWPEYGYFFQKTTNISGIIKSSEKISDCDTICQTKIGAINSTLPFQ<br>NIHQNAIGDCPKYVKAQELVLATGLRNNPIKETRGLFGAIAGFIEGGWQGLIDGWYGY<br>HHQNSEGSGYAADKEATQKAVDAITTKVNNIIDKMNTQFESTAKEFNKIEMRIKHLSDR<br>VDDGFLDVWSYNAELLVLLENERTLDFHDANVNNLYQKVKVQLKDNAIDMGNGCFKI<br>LHKCNNTCMDDIKNGTYNYYEYRKESGGRLVPRGSHHHHHH |
| H18-mut | ADPDQICIGYHSNNSTQTVNTLLESNVPVTSSHSILEKEHNGLLCKLKGKAPLDLIDCS<br>LPAWLMGNPKCDELLTASEWAYIKEDPEPENGICFPGDFDSLEDLILLVSNTDHFRKE<br>KIIDMTRFSDVTTNNVDSACPYDTNGASFYRNLNWVQQNKGKQLIFHYQNSENNPLL<br>IIWGVHQTSNAAEQNTYYGSQTGSTTITIGEETNTYPLVISESSILNGHSDRINYFWGV<br>VNPNQNFSIVSTGNFIWPEYGYFFQKTTNISGIIKSSEKISDCDTICQTKIGAINSTLPFQ<br>NIHQNAIGDCPKYVKAQELVLATGLRNNPIKETRGLFGAIAGFIEGGWQGLIDGWYGY<br>HHQNSEGSGYAADKEATQKAVDKITTKVNNIIDKMNTQFESTAKEFNKIEMRIKHLSDR<br>VDDGFLDVWSYLAELLVLLLNERTLAFHDANVNNLYQKVKVQLKDNAIDMGNGCFKIL<br>HKCNNTCMDDIKNGTYNYYEYRKESGGRLVPRGSHHHHHH |

Footnote: Non-native sequences including linkers, histidine tags and thrombin sites are written in purple.

**Supplementary Table 8 | Cryo-EM data collection statistics**

| Protein Name | H1/Cal09-WT | H1/Cal09-Wt(tri) | H1/Mal07-WT | H1/Mal07-tri |
|---|---|---|---|---|
| **Sample condition** | | | | |
| Concentration (mg/ml) | 0.6 | 0.6 | 1.0 | 1.0 |
| **Data collection and processing** | | | | |
| Micrographs | 5,330 | 3,999 | 1,078 | 847 |
| Magnification | 130,000x | 135,000x | 130,000x | 130,000x |
| Voltage(kV) | 300 | 300 | 300 | 300 |
| Electron exposure ($e^-$/Å$^2$) | 70 | 70 | 70 | 70 |
| Defocus range (μm) | -0.7 to -1.7 | -0.8 to -2.2 | -1.0 to -2.0 | -1.0 to -2.0 |
| Pixel size(Å) | 0.938 | 0.651 | 0.938 | 0.938 |
| Box size (pixels) | 288 | 288 | 256 | 256 |
| **Refinement Software** | CryoSPARC | CryoSPARC | CryoSPARC | CryoSPARC |

**Supplementary Table 9 | Cryo-EM data collection and model building statistics**

| **Protein Name** | H1/Cal09-mut | H1/Mal07-mut | H9/HK98-mut |
|---|---|---|---|
| **Sample condition** | | | |
| concentration (mg/ml) | 0.6 | 0.8 | 0.9 |
| **Data collection and processing** | | | |
| Micrographs | 3,525 | 6,246 | 5,844 |
| Magnification | 105,000x | 135,000x | 135,000x |
| Voltage(kV) | 300 | 300 | 300 |
| Electron exposure ($e^-/Å^2$) | 68.28 | 70 | 70 |
| Defocus range (μm) | -0.8 to -2.0 | -0.8 to -2.2 | -0.8 to -2.0 |
| Pixel size (Å) | 0.858 | 0.651 | 0.651 |
| Box size (pixels) | 320 | 420 | 420 |
| Symmetry imposed | C3 | C3 | C3 |
| Initial particle images (no.) | 973,699 | 426,255 | 403,550 |
| Final particle images (no.) | 464,009 | 421,214 | 430,639 |
| Map resolution (Å) | 2.22 | 2.32 | 2.27 |
| FSC threshold | 0.143 | 0.143 | 0.143 |
| **Refinement Software** | CryoSPARC | CryoSPARC | CryoSPARC |
| **Refinement** | | | |
| **Initial model used** | AlphaFold 2 | AlphaFold 3 | AlphaFold 3 |
| **Model resolution (Å)** | 2.20 | 3.00 | 2.90 |
| FSC threshold | 0.50 | 0.50 | 0.50 |
| **Map sharpening B factor ($Å^2$)** | | | |
| Protien | 31.04 | 54.20 | 48.72 |
| Glycan residues | 38.17 | - | 47.52 |
| Nucleotides | - | - | - |
| **R.m.s deviations** | | | |
| Bond length(Å) | 0.002 | 0.003 | 0.004 |
| Bond angles (˚) | 0.532 | 0.606 | 0.611 |
| **Model composition** | | | |
| Non-hydrogen atoms | 11,017 | 11,508 | 11,304 |
| Protein residues | 1,398 | 1,446 | 1,431 |
| **Validation** | | | |
| Mol Probity score | 1.14 | 1.53 | 1.61 |
| Clashscore | 2.21 | 6.91 | 8.57 |
| Poor rotamers (%) | 0.00 | 0.78 | 0.80 |
| **Ramanchandran plot** | | | |
| Favored (%) | 97.26 | 97.20 | 97.18 |
| Allowed (%) | 2.74 | 2.80 | 2.82 |
| Disallowed (%) | 0.00 | 0.00 | 0.00 |
